# Chiral shift toward D-serine reflects intrathecal inflammation in multiple sclerosis and counteracts motor impairment in a murine model

**DOI:** 10.1101/2025.05.07.652561

**Authors:** Alessandro Usiello, Kenta Arisumi, Tommaso Nuzzo, Luana Gilio, Sakiko Taniguchi, Rosita Russo, Haruhiko Motegi, Raffaella di Vito, Junichi Hata, Francesco Errico, Akinori Hashiguchi, Hideyuki Okano, Roberto Furlan, Annamaria Finardi, Mario Stampanoni Bassi, Jin Nakahara, Masashi Mita, Takanori Kanai, Angela Chambery, Masato Yasui, Diego Centonze, Jumpei Sasabe

**Affiliations:** Department of Environmental, Biological and Pharmaceutical Sciences and Technologies, University of Campania, “L. Vanvitelli”; Caserta, Italy; CEINGE Biotecnologie Avanzate “Franco Salvatore”; Naples, Italy; Department of Pharmacology, Keio University School of Medicine; Tokyo, Japan; Unit of Neurology, IRCCS Neuromed; Pozzilli (IS), Italy; Faculty of Psychology, International Telematic University UNINETTUNO; Rome, Italy; Department of Internal Medicine, Keio University School of Medicine; Tokyo, Japan; Graduate School of Human Health Sciences, Tokyo Metropolitan University; Tokyo, Japan; Keio University Regenerative Medicine Research Center; Tokyo, Japan; Laboratory for Marmoset Models of Brain Disease, RIKEN Center for Brain Science; Tokyo, Japan; Department of Agricultural Sciences, University of Naples “Federico II”; Portici, Italy; Department of Pathology, Keio University School of Medicine; Tokyo, Japan; Clinical Neuroimmunology Unit, Institute of Experimental Neurology (INSpe), Division of Neuroscience, IRCCS San Raffaele Scientific Institute, Via Olgettina, Milan, Italy; Faculty of Medicine and Surgery, Vita e Salute San Raffaele University, Via Olgettina, Milan, Italy; KAGAMI Inc.; Osaka, Japan; Department of Systems Medicine, Tor Vergata University; Rome, Italy; Laboratory of Electron Microscope and Chiral Medical Biology, Keio University; Tokyo, Japan; Human Biology-Microbiome-Quantum Research Center (WPI-Bio2Q), Keio University; Tokyo, Japan

**Author notes:** Corresponding authors. Email: Jumpei Sasabe,; Alessandro Usiello. These authors contributed equally to this work (Alessandro Usiello, Kenta Arisumi).

## Abstract

Multiple sclerosis (MS) is characterized by chronic inflammatory demyelination involving complex interplay between the central nervous and immune systems. Neuroinflammation triggers cellular reorganization requiring l-serine for sustained syntheses of membrane lipids and nucleic acids, whereas it causes aberrant glutamatergic neurotransmission involving d-serine. However, significance of serine metabolism in MS pathology remains unexplored. Here we show that serine chiral homeostasis is disrupted in MS and endogenous d-serine prevents motor deficits caused by inflammatory demyelination. We found in a large cohort study that patients with MS exhibit elevated d-serine levels and the d-/total serine ratio in the cerebrospinal fluid at diagnosis. Steric deviation toward d-serine accords with emergence of the intrathecal inflammatory marker oligoclonal bands, and correlates negatively with proinflammatory cytokines. An *in vivo* animal model of MS, genetically engineered to exhibit distinct metabolic states of d-serine, revealed that endogenous d-serine synthesis mitigates the progression of motor deficits and suppresses proinflammatory and vascular endothelial pathogenic signaling. Moreover, pre-symptomatic oral supplementation with d-serine, but not l-serine, enhances production of extracellular matrices, preserves integrity of the blood brain barrier, attenuates demyelination, and improves motor function. Contrary to the previously recognized neurotoxic nature of d-serine, our findings reveal an unrecognized significance of d-serine metabolism in MS and a protective function of d-serine against neuroinflammation involving disruption of the blood brain barrier, which may present an untapped therapeutic target in MS.

**One Sentence Summary:** Serine chiral homeostasis is disturbed in multiple sclerosis and d-serine mitigates inflammatory demyelination.

## INTRODUCTION

Multiple sclerosis (MS) is the most common inflammatory disease of the central nervous system (CNS), characterized by the sclerosis of tissues that occur both temporally and spatially. Lesions appear throughout the CNS, affecting both gray and white matter, and are associated with inflammatory demyelination, gliosis, and neurodegeneration (*1*). Focal infiltration of lymphocytes in the meninges and perivascular spaces leads to the release of soluble factors, which are thought to induce demyelination or neurodegeneration, either directly or indirectly, by activating glial cells and macrophages (*1, 2*). Therefore, tissue damage in MS results from a complex interplay between the immune system, glial cells, and neurons under inflammatory conditions. While inflammation plays a central role in tissue damage at all stages of MS pathology, metabolism that regulates MS inflammation remains less understood.

Serine metabolism serves critical functions in the architecture and operations of the CNS and immune system. l-serine is synthesized *de novo*, primarily from an intermediate of the glycolytic pathway, and is incorporated into membrane lipids to form integral components of plasma/organelle membranes and myelin (*3, 4*). Furthermore, l-serine metabolism provides one-carbon units for nucleic acid synthesis, methylation, and reductive metabolism. Since these pathways essentially support proliferative cells, undifferentiated or immune cells are susceptible to deprivation of l-serine (*5-7*). Indeed, metabolic disorders of l-serine cause congenital neural disorders with morphological abnormalities including microcephaly and hypomyelination(*8-10*). Moreover, l-serine is required for lymphocyte/macrophage proliferation and cytokine synthesis in the immune system (*6, 7*). On the other hand, l-serine uniquely undergoes functional expansion through stereo-conversion and works as a neurotransmitter precursor in the CNS. d-serine, synthesized by endogenous isomerization from l-serine (*11*), binds to *N*-methyl-d-aspartate receptors (NMDARs) with l-glutamate (*12-14*). The modulatory role of d-serine on NMDARs extends across diverse aspects of neurophysiology and contributes to the etiology and/or pathology of acquired neurological and psychiatric disorders with neuroexcitatory abnormalities (*15-27*). Notably, more recent studies indicate an anti-inflammatory role of d-serine in the periphery. d-serine suppresses the progression of chronic colitis with reduced T cell infiltration into lamina propria (*28*), and also inhibits interferon-*γ* production by inactivating mTORC1 in CD8+ T cells in the pathology of tuberculosis infection (*29*). Despite these essential roles of serine metabolism in the CNS and immune system, its significance in neuroinflammation remains largely unexplored.

## RESULTS

### Chiral equilibrium of serine shifts to the d-enantiomer in multiple sclerosis

To assess alterations of serine metabolism in MS, we first detected levels of serine enantiomers in the cerebrospinal fluid (CSF) drawn at diagnosis from patients with MS (n = 179) and other neurological disorders (OND, n = 40) (table S1). Analyses with high performance liquid chromatography (HPLC) showed that patients with MS had CSF d-serine levels elevated by 25%, with a tendency to increase l-enantiomer levels as well (Fig. 1A-C), resulting in 13% increase in the d/total-serine ratio (Fig. 1D), compared with patients with OND. Notably, among patients with MS, the enhanced d/total-serine ratio was associated with emergence of IgG oligoclonal bands (OCBs) in the CSF (Fig. 1E and fig. S1AB), a hallmark of MS (*30*), suggesting that serine chirality reflects a CSF-restricted immune response in the CNS.

**Fig. 1.**
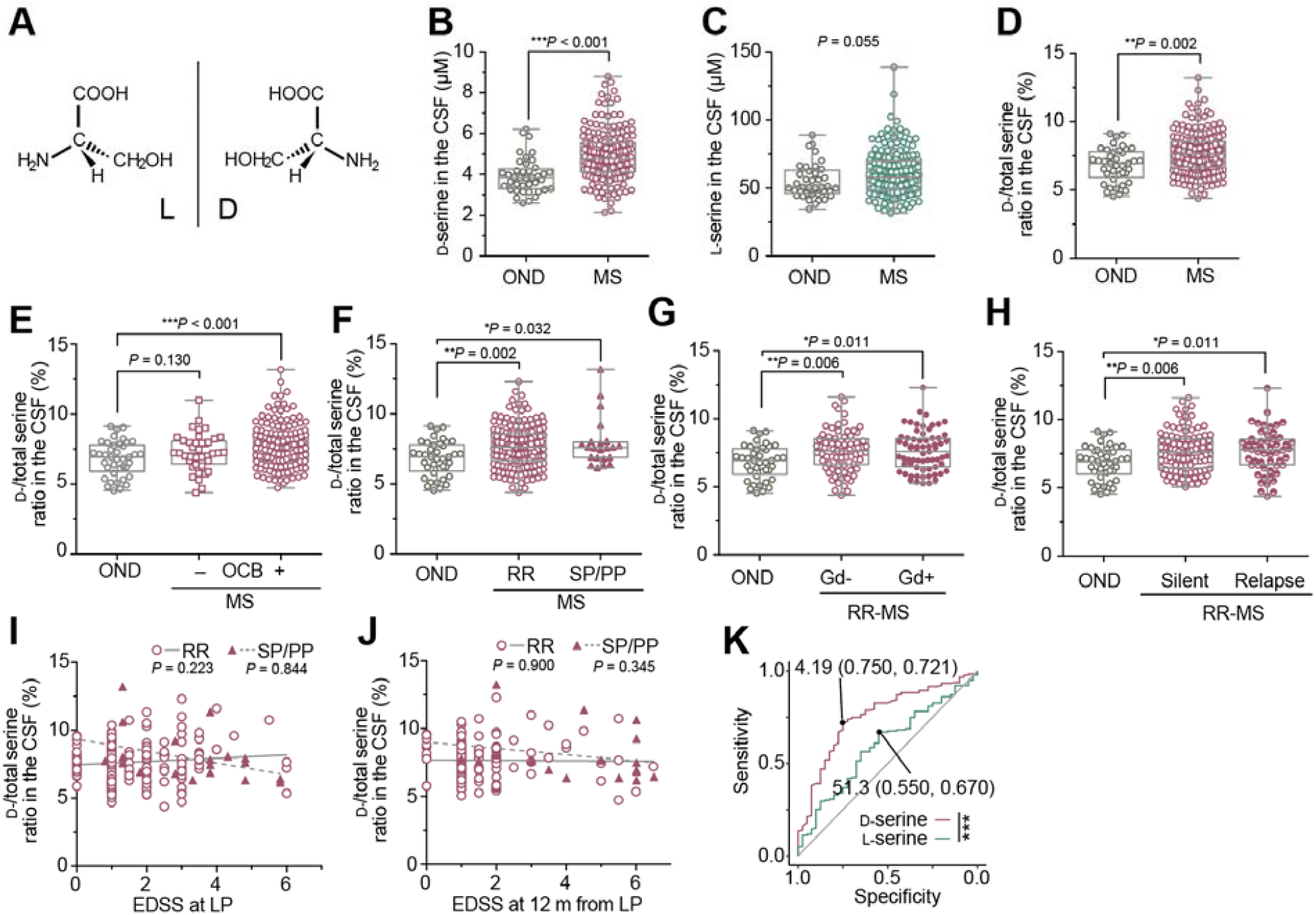
d-enantiomer-dominant increase of serine in CSF of patients with MS. CSF wa drawn from patients with OND (n = 40) and MS (n = 179) at diagnosis. (**A**) Ser enantiomers were measured with HPLC. (**B** and **C**) CSF levels of D- (B) or L-serine (C) in patients with OND and MS. (**D**-**H**) D-/total serine ratio in the CSF of OND and whole subjects of MS (D); MS subjects categorized with detectability of OCB (E) or disease types (RR and SP/PP) (F); or subjects with RR-MS categorized with Gd extravasation in MRI (G) or disease activity (H). (**I** and **J**) Correlations between D-/total serine ratio and EDSS scores in patients with RR- or SP/PP-MS at lumbar puncture (LP) (I) or at 12 months after LP (J). (**K**) True/false positive rates to detect MS at each threshold setting for D- or L-serine. Values show cut-offs (µM) with (specificity, sensitivity). Data ar shown in box plots (b-h). \*\*\**P* < 0.001, \*\**P* < 0.01, \**P* < 0.05; Mann-Whitney test (B-H), linear regression analysis (I and J), and DeLong’s test (K).

On the other hand, d-serine levels or d/total-serine ratios in the CSF at diagnosis did not predict the disease course of MS (relapsing-remitting, RR; secondary progressive, SP; and primary progressive, PP) based on revised McDonald criteria (*31*)(Fig. 1F and fig. S1C). Moreover, these parameters did not reflect extravasation of gadolinium (Gd) on MRI (Fig. 1G and fig. S1E), disease phases in RR-MS patients (Fig. 1H and fig. S1G), or clinical scores of disability, quantified using the Expanded Disability Status Scale (EDSS) at diagnosis or 12 months later (Fig. 1IJ and fig. S2). Therefore, d-serine accumulation in the CSF at diagnosis does not represent disease activity of MS, but appears to be one of its common neurochemical features. In contrast, an l-serine increase was evident only in RR-MS patients (fig. S1D), especially those lacking Gd-enhancing lesions on MRI or in the silent phase (fig. S1FH), implying that an l-serine increase is a sign of inactive disease, rather than a general characteristic of MS. Indeed, receiver operator characteristic (ROC) curves for serine enantiomers in the CSF show that d-serine detects MS more specifically and sensitively than l-serine (Fig. 1K) (cut-off values for d-serine, 4.190 µM; for l-serine, 51.32 µM).

### CSF d-serine shows linear correlations with cytokines in MS pathology

The nervous and immune systems have mutual interactions that modulate tissue homeostasis and inflammation (*32*). Consistent with expression of diverse functional ionotropic/metabotropic glutamate receptors, including NMDARs, in the nervous and immune systems (*33*), l-glutamate in CSF of patients with MS is associated with neuroinflammation (*34, 35*). Given the requirement of d-serine with l-glutamate in activation of NMDARs, we wondered whether variations of d-serine are linked to CNS immune responses in MS. Spearman’s rank correlation coefficient analyses showed that among cytokines / chemokines / growth factors, in CSF of MS patients, d-serine had mild negative correlations with some proinflammatory cytokines, such as IL-6 and IL-15. Conversely, it was positively correlated with an anti-inflammatory IL-1 receptor antagonist (IL-1ra) (Fig. 2A). Regression analyses indicated that such correlations had linear associations between d-serine and pro-/anti-inflammatory cytokines in CSF from patients with MS (Fig. 2B-G and fig. S3). Intriguingly, whereas d-serine had a positive linear association with l-serine in the CSF from both OND and MS patients (fig. S4AB), l-glutamate showed a clear negative association with d-serine only in patients with MS (fig. S4CD and 5)(*34*). Given that enhanced excitatory signaling by high levels of l-glutamate triggers degeneration of neurons and oligodendrocytes in MS patients and animal models (*33, 36, 37*), our results showing negative correlations between d-serine and proinflammatory signals in the CSF of MS patients suggest that d-serine exerts neuroprotective effects, opposing those of l-glutamate.

**Fig. 2.**
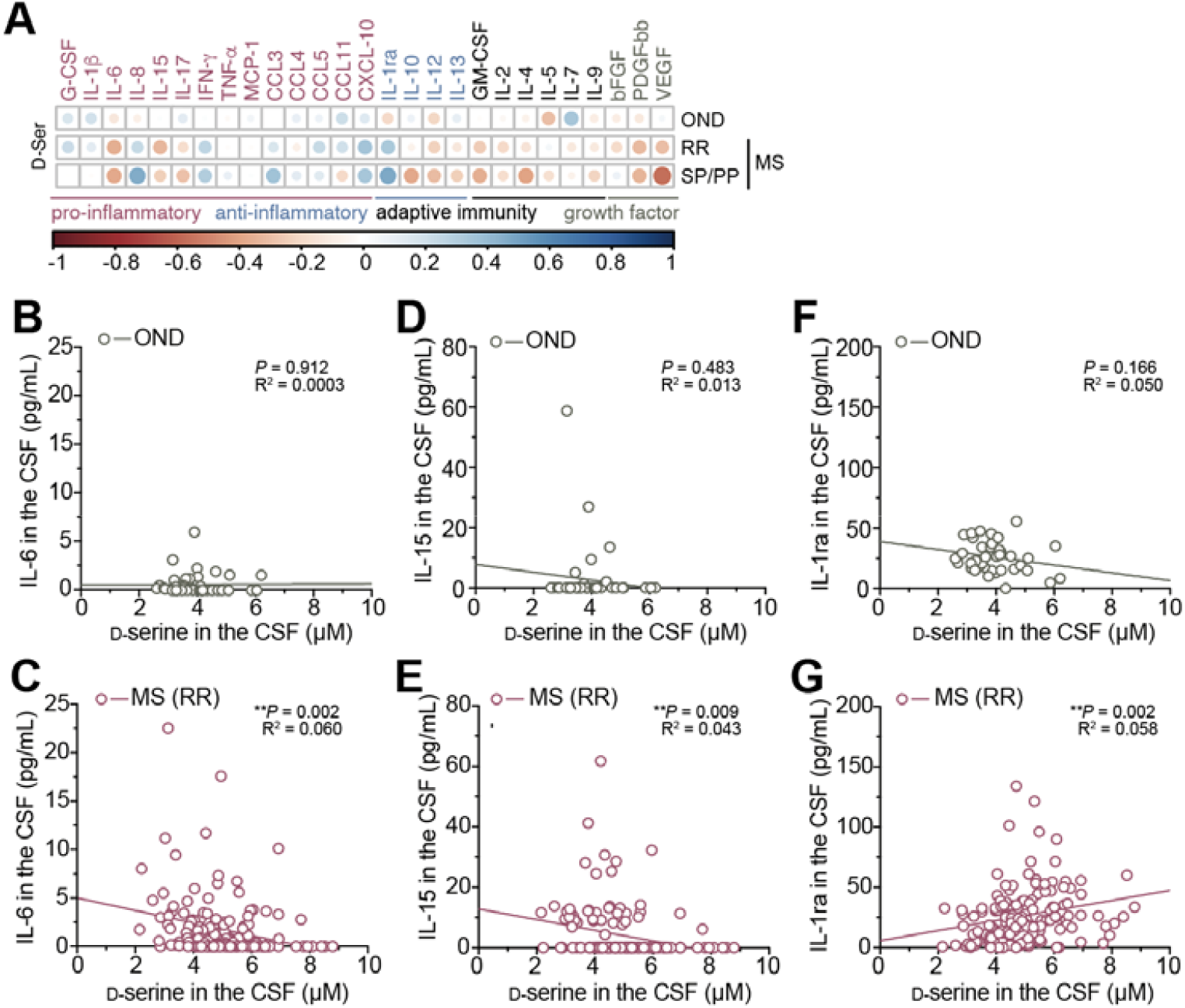
Associations of d-serine with inflammatory cytokines in CSF with MS. (**A**) A heatmap showing Spearman’s correlation coefficients between D-serine and cytokine levels in CSF from patients with OND (n = 40), RR-MS (n = 157), and SP/PP-MS (n = 22). (**B**-**G**) Correlations between D-serine and IL-6 (B and C), IL-15 (D and E), and IL-1ra (F and G) in CSF with OND (B, D, and F) and RR-MS (C, E, and G). \*\**P* < 0.01; linear regression.

### EAE triggers aberrant metabolism of serine enantiomers

In the CNS, l-serine is primarily synthesized in astrocytes from an intermediate of glycolysis through the phosphorylated pathway, in which phosphoglycerate dehydrogenase (Phgdh) works as the rate-limiting enzyme (*38*). l-serine, shuttled from astrocytes, is chiral-converted by serine racemase (Sr) in neurons (*39, 40*). Released d-serine is then taken up by astrocytes and degraded by d-amino acid oxidase (Dao)(*41*) (Fig. 3A). To further examine serine homeostasis in neuroinflammation, we used mice sensitized to peptide fragment 35-55 of myelin oligodendrocyte glycoprotein (MOG_35-55_) as an animal model of MS (Fig. 3B). Levels of both serine enantiomers were unaltered in spinal cord with experimental autoimmune encephalomyelitis (EAE) at a pre-symptomatic phase (day 7 after MOG_35-55_ induction), but significantly increased at a chronic phase (day 30) (Fig. 3CD), indicating that active neuroinflammation is associated with serine accumulation. d/total-serine ratios in EAE spinal cord remained unchanged compared to those in non-EAE controls (Fig. 3E), suggesting that the chiral-conversion rate of serine is not affected in EAE. In accordance with the increase of l-serine under EAE, transcription and expression of Phgdh were significantly elevated in the spinal cord, but not in the cerebral cortex or cerebellum (fig. S6A), at an acute phase (Fig. 3FG). On the other hand, expression of Dao and Sr showed no detectable alteration, despite a striking reduction of Dao transcription (Fig. 3FG and fig. S6B). Since we observed significant accumulation of d-serine, we further studied whether degradation of d-serine is also reduced in the EAE spinal cord. A challenge of d-serine in the drinking water resulted in strikingly enhanced accumulation of d-serine with no alteration of l-serine in EAE spinal cord at a chronic, but not pre-symptomatic, phase (Fig. 3HI), indicating that in addition to enhanced synthesis of l-serine, degradation of the d-enantiomer is disturbed in the CNS under EAE. Since Phgdh and Dao are both astrocytic enzymes (Fig. 3A), these results suggest that pathologic astrocytes produce excessive l-serine and disturb degradation of d-serine under demyelinating and neurodegenerative processes typically associated with EAE.

**Fig. 3.**
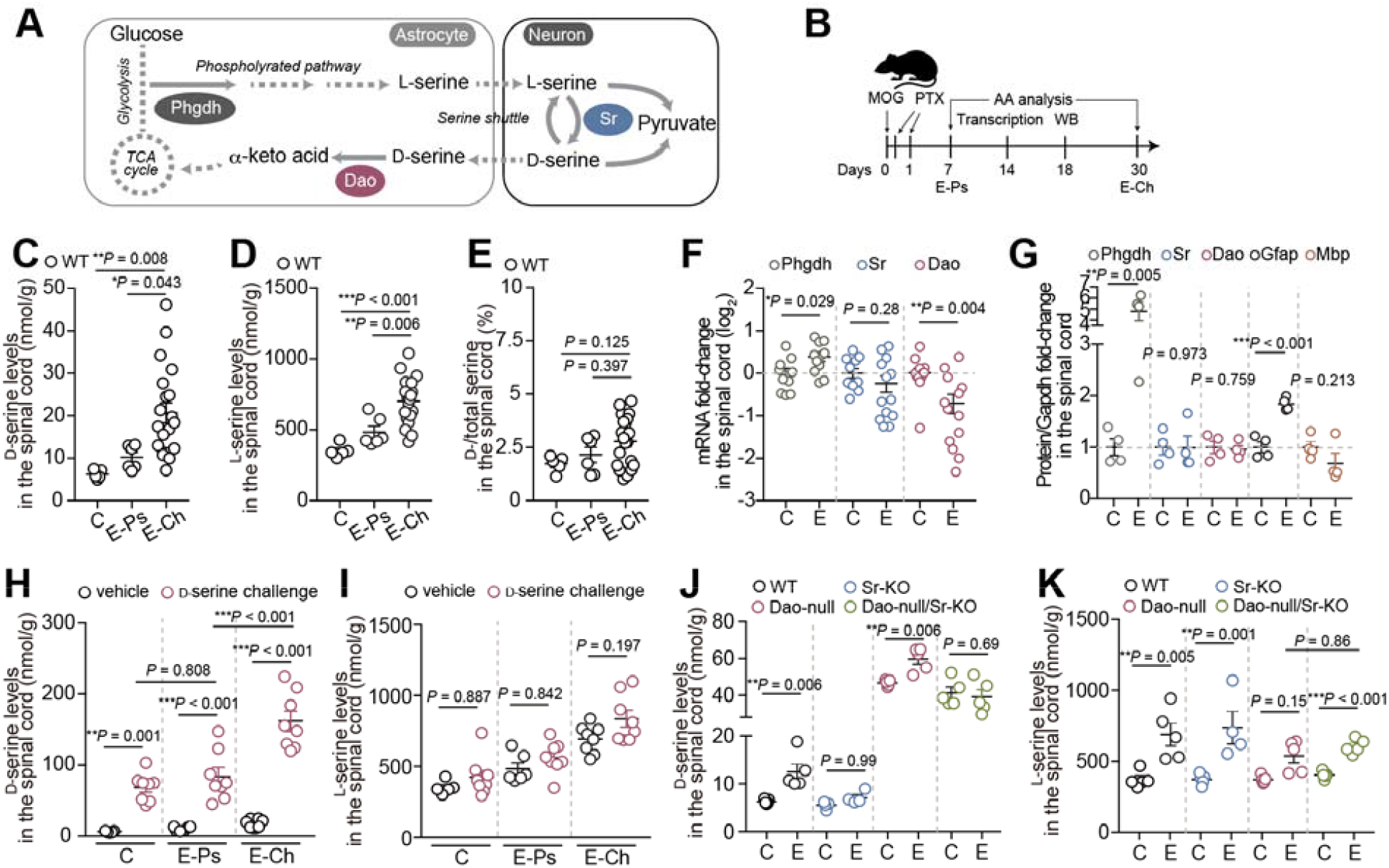
EAE disturbs homeostasis of serine enantiomers. (**A**) Major metabolism of serine enantiomers. **b**, Sampling schedule after EAE induction. AA, amino acids. (**C**-**E** and **H**-**K**) D- (C, H, and J) / L-serine levels (D, I, and K) or D-/total serine ratio (E) in spinal cords from wild-type (WT) (C-E and H-K), Sr knockout (Sr-KO) (J and K), Dao-null (and K), Dao-null/Sr-KO (J and K) mice with or without EAE. For wild-type mice, D-serine (30 mM in the drinking water) was further challenged for 1 week (H and I). (**F** and **G**) mRNA (F) and protein (G) expression of serine metabolic enzymes standardized against Gapdh in spinal cords from EAE and control mice. Data are shown as means ± s.e.m. (C-K). C, control. E, EAE. E-Ps/Ch, pre-symptomatic/chronic stages of EAE. \*\*\**P* < 0.001, \*\**P* < 0.01, \**P* < 0.05; one-way ANOVA with Tukey’s test (C-E and H-K) and unpaired two-tailed t-test (F and G).

### Endogenous d-serine synthesis suppresses proinflammatory and vascular remodeling signals, and ameliorates motor deficits in EAE

We then evaluated contributions of Dao and Sr to d-serine metabolism and disease progression in EAE animals. Genetic loss of Sr diminished the EAE-induced increase of d-serine, whereas Dao deficiency due to a G181R-point mutation (a null-mutation) enhanced the increase of d-serine in EAE spinal cord (Fig. 3J). Elevated d-serine in Dao-null mice with EAE was significantly reduced in double Dao-null/Sr-knockout mice with EAE (Fig. 3J), showing that the EAE-induced increase of d-serine essentially mediates chiral conversion by Sr. On the other hand, l-serine levels were not altered under EAE by loss of Sr, Dao, or both (Fig. 3K), supporting observations that Sr and Dao catalyze reactions enantio-selectively to the d-configuration of serine. Notably, motor disability in EAE was strictly associated with basal levels of d-serine in mouse spinal cord. Indeed, lack of d-serine synthesis by Sr knockout caused a dramatic exacerbation of motor deficits in EAE, although EAE onset was not altered (Fig. 4AB). On the contrary, enzymatic inactivation of Dao, giving rise to a baseline accumulation of d-serine, prominently attenuated motor symptoms of EAE mice (Fig. 4B). The remarkable improvement of motor function by Dao deficiency was nullified by loss of d-serine synthesis due to additional genetic disruption of Sr (Fig. 4C). Although Dao also degrades diverse d-amino acids originating from gut microbes and modulates intestinal immune responses(*42, 43*), gut-conditional knockout of Dao did not impact the disease course of EAE (Fig. 4D). Therefore, those results show that metabolism of endogenous d-serine, but not microbial d-amino acids is a crucial determinant of EAE progression in mice. Consistent with changes of motor impairment in different metabolic states of d-serine under EAE, *q*-space diffusion MRI, a technique that visualizes myelin, showed significant attenuation of EAE-induced demyelination in Dao-null mice compared to wild-type or Sr-knockout mice (Fig. 4E and fig. S7).

**Fig. 4.**
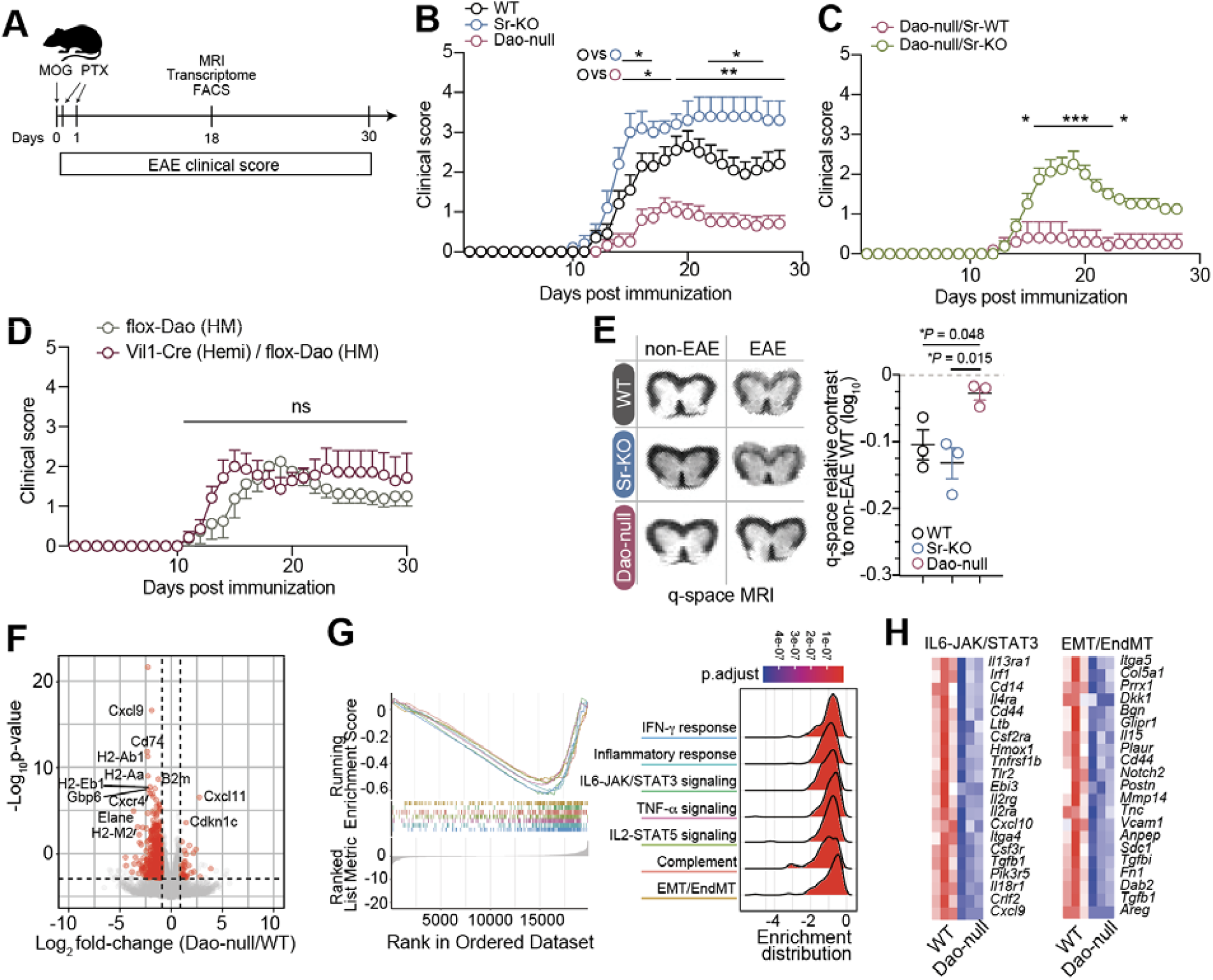
d-serine metabolism is a major determinant of EAE progression. (**A**) Experimental schedule after EAE induction. (**B**-**D**) Clinical scores after MOG injection for WT, Sr-KO, and Dao-null mice (n = 5, 10, 10) (B); Dao-null/Sr-WT and Dao-null/Sr-KO mice (n = 8, 5) (C); and flox-Dao and Vil1-Cre/flox-Dao mice (n = 8, 7) (D). (**E**) Myelin imaging by *q*-space diffusion MRI for spinal cords from WT, Sr-KO, and Dao-null mice with or without EAE. Representative images (left). Relative contrast in the white matter of mic with EAE to that of non-EAE WT controls (right). Data are shown as means ± s.e.m. (**F**-**H**) RNA-seq of spinal cords from Dao-null and WT mice. A volcano plot indicating differentially expressed genes in Dao-null mice compared to WT (F). GSEA plots and Ridgeplots for a core subset of gene ontologies mRNA (G). Heatmaps showing z-scores of gene expression significantly altered by Dao (H). Color represents fold-change in expression in Dao-null mice compared to WT. \*\*\**P* < 0.001, \*\**P* < 0.01, \**P* < 0.05; two-way ANOVA with Sidak’s test (B-D) and one-way ANOVA with Tukey’s test (E).

To further understand attenuation of the neuroinflammatory phenotype in Dao-null mice at molecular levels, we performed RNA-sequencing using spinal cords of Dao-null and wild-type mice at an acute phase of EAE. Principal component analysis (PCA) and a volcano plot of the log_2_ fold-change in the gene expression profile showed that Dao activity altered transcriptional activity strikingly in EAE (Fig. 4F and fig. S8a). Gene set enrichment analysis (GSEA) using the Molecular Signatures Database (MSigDB) revealed that lack of Dao activity reduced transcription of diverse gene sets associated with complement activation, responses to inflammation and IFN-*γ*, as well as signaling pathways for IL6-JAK/STAT3, TNF-*α*, and IL2-STAT5 (Fig. 4GH and fig. S8B). Such inhibitory actions against proinflammatory pathways by loss of Dao were also confirmed by GSEA of gene ontology using the ClusterProfiler package (fig. S8C). Notably, in addition to the anti-inflammatory transcriptional profile, we observed a striking decrease in transcription of gene sets, associated with TGF-*β* signaling, Notch pathway, and degradation of matrix proteins, for epithelial mesenchymal transition (EMT) and endothelial-to-mesenchymal transition (EndMT) in DAO-null spinal cords (Fig. 4GH). EndMT is a pathophysiological process in which endothelial cells progressively transdifferentiate into mesenchymal cells and lose endothelial features. Since crosstalk between proinflammatory and EndMT signaling occurs before onset and initiates breakdown of the blood-brain barrier (BBB) in EAE mice (*44*), our findings indicate that loss of d-serine degradation attenuates both proinflammatory pathways and processes of vascular endothelial pathology.

### Exogenous d-serine preserves BBB integrity and improves motor function in EAE

A growing body of evidence suggests that disruption of the BBB serves a critical pathogenic function in initial progression and disease amplification in MS (*45, 46*). Given the transcriptional profiles associated with EndMT in Dao-null mice (Fig. 4H), we hypothesized that d-serine intake might counter BBB disruption in the EAE model. To test this hypothesis, we treated C57BL6 mice with d-serine in the drinking water before and after MOG sensitization (Fig. 5A). The concentration of d-serine in the drinking water was determined so that d-serine levels in the spinal cord were equivalent to those of Dao-null mice (fig. S9A-D)(*47*). We studied quantitative proteome profiling at a pre-symptomatic stage of EAE under treatment with d-serine or vehicle, and identified 4,900 high-confidence proteins on average in these spinal cords (Fig. 5B and fig. S10A). Of these, d-serine treatment altered expression of 47 proteins in spinal cords (fig. S10B). Gene ontology enrichment analysis for differentially expressed proteins revealed that d-serine enhanced molecular functions related to extracellular matrix structure binding and constituent-related terms, calcium-dependent phospholipid and protein binding, and fatty acid binding and other lipid-related functions (Fig. 5BC and fig. S10BC). Changes in expression of these extracellular and membrane-associated proteins, including collagen alpha (I/VI), annexin A1/2/4, caveolin-1, transgelin, fibrillin, and biglycan, in the spinal cord before onset of EAE suggested that d-serine impacts stromal function such as basement membrane integrity. Indeed, d-serine treatment for two weeks enhanced protein expression of annexin A2 (fig. S10D), which modulates BBB permeability (*48*), and significantly inhibited the EAE-induced increase of BBB transparency, especially in spinal cords at onset of the disease (Fig. 5D). Electron microscopy of white matter in the spinal cord with EAE shows that d-serine treatment preserved integrity of basement membrane surrounding micro-vessels as well as myelin structure (Fig. 5E and fig. S11). Furthermore, in accordance with the attenuated disease severity of EAE in Dao-null mice compared to wild-type, d-serine intake before and after MOG injection throughout the course conspicuously suppressed progression of EAE, whereas l-enantiomer treatment did not impact the disease course (Fig. 5F). This *in vivo* protective effect of d-serine against EAE was enantio-selective, as d-serine intake did not alter l-serine levels in spinal cords of EAE mice, and vice versa (fig. S12A). We further tested whether timing of d-serine treatment influences its protective effect (fig. S12B). A protective effect against EAE was observed even when d-serine intake was terminated 7 days after MOG sensitization (fig. S12C), whereas there was no effect when d-serine treatment was started on the day of MOG injection and continued thereafter (fig. S12D). d-serine concentrations in spinal cords at a chronic phase in these two conditions were not associated with the protective effect of d-serine on EAE (fig. S12EF), indicating that maintaining high d-serine levels early in the disease is crucial in attenuating EAE progression. Thus, our results suggest that increased d-serine availability in the CNS impacts initial development of EAE by modulating stromal functions and permeability of the BBB.

**Fig. 5.**
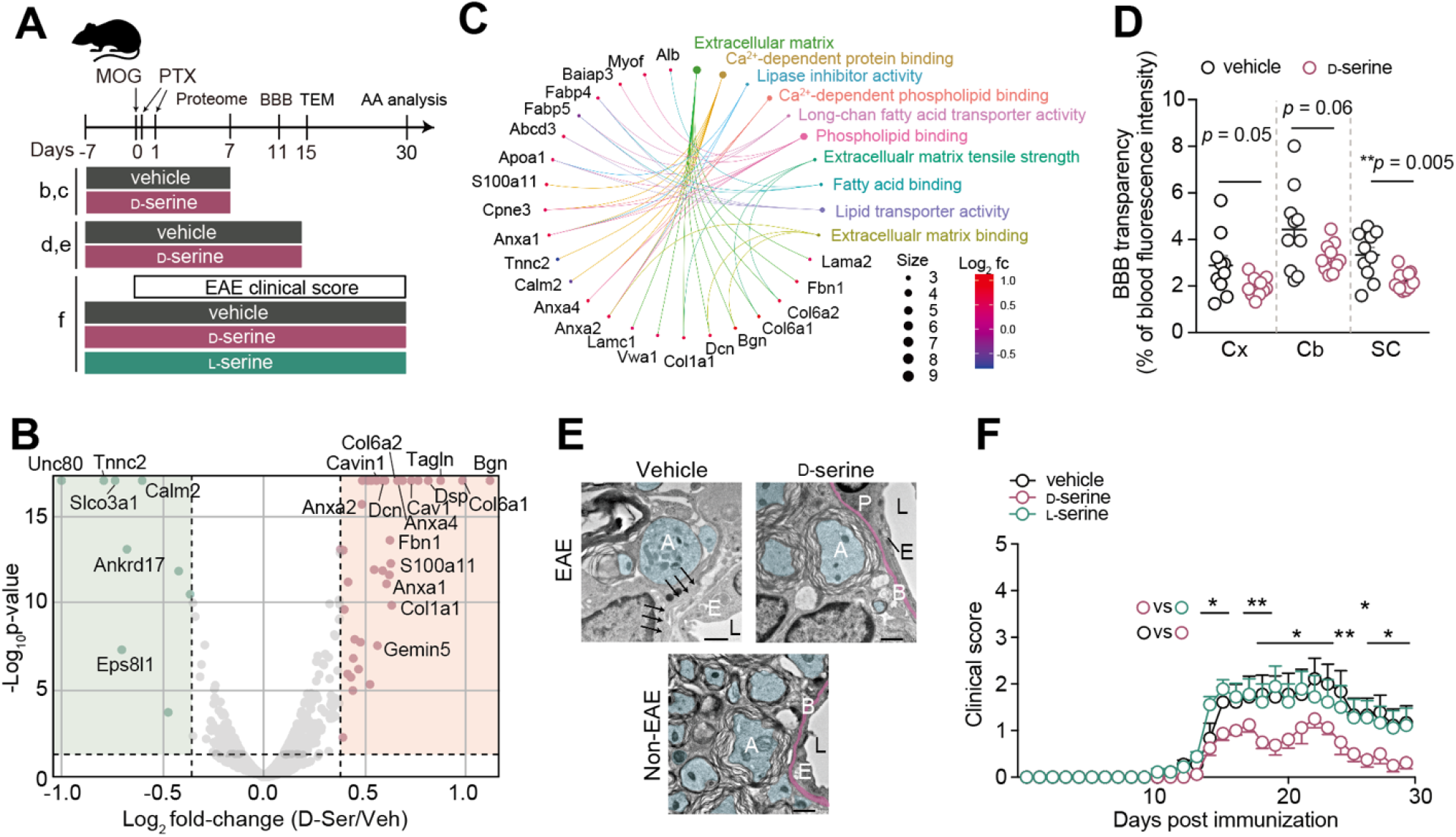
d-serine improves BBB integrity and hinders EAE progression. (**A**) Experimental schedule designed for (B-F). TEM, transmission electron microscopy. (**B** and **C**) Proteomic analysis of spinal cords from wild-type mice treated with D-serine in their drinking water or vehicle. A volcano plot indicating differentially expressed proteins (DEPs) in D-serine-treated mice compared to controls (B). A CNET plot showing the strength of association between DEPs and representative GO enrichment terms (C). (**D**) Transparency of the BBB in cerebral cortex (Cx), cerebellum (Cb), and spinal cord (SC) examined using D-serine- or vehicle-treated mice injected with fluorescein-dextran (n = 10 each). (**E**) Representative TEM images of the anterior funiculus in spinal cords from D-serine- or vehicle-treated EAE mice or non-EAE control. Scale bars, 1 µm. A, axon (blue). B, basement membrane (magenta). E, vascular endothelial cell. L, vascular lumen. Arrows, disrupted basement membrane. (**F**) Clinical scores after MOG injection for WT mice treated with D- or L-serine or vehicle in the drinking water (n = 9, 9, 8). Data are shown as mean ± s.e.m. (D and F). Serine concentrations in the drinking water, 30 mM (B-F). \*\**P* < 0.01, \**P* < 0.05; Mann-Whitney test (D) and two-way ANOVA with Sidak’s test (F).

## DISCUSSION

Demyelination in MS is accompanied by a cascade of multiple events: sensitization and activation of peripheral lymphocytes, increased permeability of the BBB, and infiltration of lymphocytes into the CNS (*37*). Accumulating evidence indicates that disruption of the BBB is a critical pathogenic step in initiation and amplification of neuroinflammation in MS (*44-46*). Even though inflammatory stimuli promote enhanced BBB permeability through microvasculature reorganization, the intrinsic mechanism that protects against BBB disruption remains unknown. Given the crucial roles of serine metabolism in structure, growth, and function in the CNS, we used multimodal techniques to comprehensively understand the contribution of serine metabolism to MS pathogenesis using well-characterized large clinical samples and *in vivo* models. Here we show that chiral equilibrium of serine is shifted to the d-enantiomer in patients with MS and stereo-conversion from l-into d-serine significantly attenuates development of neuroinflammation, demyelination, and motor deficits in EAE mice in part by modulating BBB permeability. These findings suggest that d-serine may serve not only as a biochemical marker of intrathecal inflammation but also as a functional component of the neuroimmune response in MS.

In our large cohort of patients newly diagnosed as MS, CSF d-serine levels and the d-/total serine ratio were significantly elevated, and this increase was closely associated with the detectability of OCBs (Fig. 1BDE). Notably, these changes at diagnosis were independent of radiological phenotype, clinical disease activity, or disability scores, suggesting that this enantiomeric shift reflects early neuroinflammatory processes that may precede overt disease progression (Fig. 1E-J). On the other hand, l-serine levels were elevated especially in patients with RR-MS during clinically stable phases (fig. S1), indicating a potential role in immunological homeostasis or tissue repair processes including remyelination. These enantiomer-specific alterations in CSF support the hypothesis that dysregulation of serine chiral equilibrium is an early and mechanistically relevant feature of MS. Serine is a rare amino acid in that both its l- and d-enantiomers are biologically active in the CNS. Despite their distinct roles, prior studies on MS have not differentiated between these two forms (*49, 50*). Most untargeted metabolomic analyses do not separate the enantiomers (*51-53*), which may have led to an underestimation of enantiomer-specific changes in the MS pathology. While CSF d-serine alterations have been described in other neuropsychiatric conditions (*54, 55*), our observation that its increase coincides with OCBs positivity–an established marker of intrathecal inflammation–suggests a mechanistic link. Multiple factors may contribute to altered CSF amino acid levels in MS, including changes in parenchymal metabolism, increased permeability of BBB or the blood-CSF barrier (BCSFB), and metabolic activity of infiltrating immune cells. DAO is expressed not only in astrocytes (*41*) but is reported also in the meninges and choroid plexus (*56*), a key component of BCSFB. Therefore, elevated d-serine in CSF may reflect inflammation in meningeal and periventricular regions, beyond the brain parenchyma. Given that production of OCBs in MS is driven by intrathecal immune cell activity, the parallel increase in OCBs and d-serine supports the idea that d-serine could serve as a surrogate marker for CNS inflammation in MS. Intriguingly, d-serine levels were negatively correlated with IL-6 and IL-15–cytokines implicated in MS exacerbation (*57*)–and positively correlated with IL-1ra, which is neuroprotective in both MS and EAE models (*34*)(Fig. 2). This immunological profile is particularly striking when considered alongside our prior observation that CSF l-glutamate levels positively correlate with IL-6 and inversely with IL-1ra in the same cohort (*34*). Additionally, we found an inverse relationship between d-serine and l-glutamate levels in CSF of patients with MS (fig. S4). These complementary patterns suggest a functional dichotomy: d-serine may counteract the excitotoxic and proinflammatory roles of l-glutamate in MS pathophysiology.

In the EAE mice, d-serine progressively accumulated in the spinal cord in correlation with disease severity (Fig. 3C). This accumulation was driven by both an increase in its precursor l-serine and a reduction in d-enantiomer-specific degradation (Fig. 3DHIJ). Astrocytes synthesize l-serine, which supports cellular proliferation through one-carbon metabolism and sphingolipid biosynthesis (*4, 58*). Inflammatory activation of glia and immune cells likely increases demand for l-serine due to enhanced membrane and nucleic acid synthesis, resulting in Phgdh upregulation in astrocytes (Fig. 3FG). Additionally, demyelination may further elevate l-serine demand in oligodendrocytes to support sphingomyelin production for remyelination–though whether tissue increase in l-serine reflects active synthesis or unmetabolized accumulation remains uncertain. In parallel, increased l-serine likely drives conversion to d-serine via Sr, while decreased Dao activity allows d-serine to accumulate. This pattern mirrors observations in other neurological disorders, including cerebral ischemia (*24, 25*) and amyotrophic lateral sclerosis (*22*), which involve aberrant neuroexcitation in the pathophysiology. Since d-serine is required as a co-agonist to activate NMDARs with l-glutamate, tissue accumulation of d-serine has classically been implicated in excitotoxicity in neurological disorders (*15, 21, 24*). Involvement of excitotoxicity in MS is supported by preclinical evidence showing the improvement of functional outcomes in the EAE model by blockade of glutamatergic receptors (*59, 60*). Indeed, excess release of l-glutamate is observed during early stages of progressive MS (*61*). However, in this study, d-serine did not exacerbate excitotoxicity; instead, it exerted anti-inflammatory effects and improved clinical scores in the EAE model, revealing a previously unrecognized stereoselective neuroprotective function (Figs. 4 and 5).

Unlike l-serine, d-serine has high affinity for activity-modulating subunits, GluN1 and GluN3, of NMDARs. Classically, in several neurological disorders (*15, 21, 22, 24, 25*), d-serine has been linked to neurotoxicity through binding to GluN1-mediated NMDAR activation. In contrast, our findings that d-serine inversely correlate with l-glutamate in MS (Figs. S4), and that d-serine preserves BBB integrity in EAE mice (Figs. 5), suggest that d-serine serves a function beyond neuronal excitation. Notably, NMDARs are also expressed in vascular endothelial cells, where GluN3-containing receptors exert non-ionotropic effects to modulate cerebral blood flow and BBB permeability (*62, 63*). Glycine, another co-agonist of NMDARs, activates these GluN3-containing NMDARs, whereas d-serine antagonizes this action of glycine (*64*). Given that glycine impairs BBB integrity through activation of endothelial NMDARs (*65*), the protective effect of d-serine on BBB permeability (Fig. 5D) may indicate the involvement of endothelial NMDARs in the anti-inflammatory role of d-serine. Interestingly, d-serine enhances extracellular matrix integrity around the onset of EAE (Fig. 5BC) and ameliorates clinical phenotype with preserved morphology of vascular basement membrane (Fig. 5EF). Since inflammatory BBB damage typically occurs around symptom onset in the early stages of MS and during relapses in RR-MS (*66*), these findings suggest that d-serine may hold therapeutic potential in the initial phase of MS or in preventing relapse and maintaining remission by stabilizing BBB integrity. Nevertheless, further mechanistic studies are needed to examine spatiotemporal d-serine accumulation and its impact on BBB integrity in MS pathology. In particular, since astrocytic serine metabolic enzymes, Phgdh and Dao, show disease-associated alterations (Fig. 3FG), how astrocytes exert BBB protection in MS pathology is an intriguing question, as it would directly open the possibility of a new therapeutic strategy. Given strategic localization of perivascular endfeet of astrocytes enwrapping the vasculature, it is necessary to investigate involvement of BBB constituent cells, such as endothelial cells and pericytes, in the anti-neuroinflammatory function of d-serine (for instance, to determine whether it directly influences endothelial reorganization or production of extracellular matrices). Such investigations could potentially shed light on mechanisms that contribute to d-serine-mediated inhibition of neuroinflammation.

This study has following limitations. First, the clinical data were obtained exclusively from a cohort enrolled in Italy, and the findings have not been evaluated for generalizability across diverse racial or ethnic populations. Second, CSF samples were collected at the time of diagnosis, so the utility of d-serine as a longitudinal biomarker for disease activity remains to be determined. Third, while the EAE model captures key immunopathological aspects of MS, it does not fully recapitulate the complexity of human disease (*67*). Therefore, the observed anti-inflammatory effects of d-serine in EAE mice require validation in other MS models and clinical studies.

Together, our findings identify d-serine as a stereoselective modulator of neuroinflammation with potential to preserve BBB integrity in MS. These insights not only underscore the relevance of amino acid chiral balance in CNS immune regulation but also highlight d-serine as a promising biomarker and therapeutic target in early-stage MS.

## MATERIALS AND METHODS

### Study design

Our objective was to understand whether intrathecal inflammation affects serine chiral homeostasis in patients with MS and how serine chirality impacts EAE pathology and progression in mice. In the clinical study, patients underwent comprehensive clinical evaluations including CSF withdrawal required for diagnosis and appropriate treatments with longitudinal clinical evaluations (end points: clinical characteristics, Gd-enhanced MRI, and CSF analysis of amino acids and cytokines). Patients were not assigned randomly to experimental groups. Patients with MS (n = 179) and OND (n = 40) had full evaluation and CSF withdrawal. No statistical methods were used to predetermine sample size. Inclusion criteria were the established MS diagnosis and the ability to provide written informed consent to the study. No participants were excluded from further analyses. In mice, we tested the impact of Sr/Dao genetic depletion/inactivation and d-/l-serine treatment on EAE progression (end points: motor function and survival) and molecular biological and morphological phenotype in the CNS tissues (end points: amino acid quantification, qPCR, western blotting, transcriptome/proteome analyses, MRI, and electron microscopic analysis). In vivo, we probed the relationship between D-serine and BBB transparency in EAE mice (end points: fluorescent transparency into CNS tissue). Mice with significant leakage of subcutaneously injected MOG were excluded from further analyses. No randomization or statistical predetermination of sample size was used. Exact numbers of mice are plotted in each figure or described in each legend. n = 3 to 15 mice per group were used.

### Patient enrolment and CSF collection

179 consecutive MS patients were enrolled in the study between 2017-2019(*68*). Patients were admitted to the Neurology Unit of IRCCS Neuromed in Pozzilli (IS) and later diagnosed as suffering from RR or SP/PP MS. For diagnostic purposes, all patients underwent blood tests, complete neurological evaluations, brain and spinal MRI scans, and CSF withdrawal within 24 h. Patients were medication-free before CSF withdrawal and neurophysiological assessment. Corticosteroids or other MS-specific immunosuppressive therapies were initiated later as appropriate. The control group comprised 40 patients with non-inflammatory/non-degenerative CNS disorders or peripheral nervous system disorders, such as vascular leukoencephalopathy, metabolic and hereditary polyneuropathies, normal pressure hydrocephalus, functional neurological disorder, and spondylotic myelopathy. All patients and/or their legal representatives gave informed written consent for participation in the study. The protocol was authorized by the ethics committee of IRCCS Neuromed (cod. 10-17), and all procedures were performed in accordance with approved guidelines.

Diagnosis of RR-MS or SP/PP-MS was established by clinical, laboratory, and MRI parameters, and matched published criteria (*31*). Demographic and clinical information was derived from medical records. MS disease onset was defined as the first episode of focal neurological dysfunction indicative of MS. Disease duration was estimated as the number of months from onset to the time of diagnosis. Disability was determined by a specially trained and certified examining neurologist (Neurostatus training and documentation DVD for a standardized neurological examination and assessment of Kurtzke’s functional systems and Expanded Disability Status Scale for MS patients. Basel, Switzerland: Neurostatus, 2006; available at http://www.neurostatus.net), using the Expanded Disability Status Scale (EDSS)(*69*).

MRI examination consisted of 3 Tesla dual-echo proton density, fluid-attenuated inversion recovery (FLAIR), T2-weighted, spin-echo images and pre-contrast and post-contrast T1-weighted spin-echo images. All images were acquired in the axial orientation with 3-mm contiguous slices. Radiological activity was defined as the presence of gadolinium (Gd) (0.2 mL/Kg e.v.) enhancing lesions evaluated by a neuroradiologist who was unaware of patient clinical details. CSF samples were collected according to international guidelines (*70*). Lumbar puncture (LP) was performed from AM 8:00 to 10:00, after overnight fasting. Portions (2 mL) of CSF samples were used for biochemical analyses. CSF was immediately collected in sterile polypropylene tubes (Sarstedt® tubes, codes: 62.610.210) and gently mixed to avoid possible gradient effects. All samples were centrifuged at 2,000 x g for 10 min at room temperature and aliquoted in 0.5-mL portions into sterile polypropylene tubes (Sarstedt® tubes, codes: 72.730.007). Aliquots were stored at −80°C until use, avoiding freeze/thaw cycles. Blood-contaminated samples were excluded from the analysis (cut-off of 50 red blood cells per µL). Internal quality controls were assayed in each run. Operators blinded to the diagnosis performed these measurements.

### Quantification of serine enantiomers

Human CSF samples were analysed for serine enantiomers as previously reported(*26*). Clinical CSF samples (100 µL) were mixed in a 1:10 dilution with HPLC-grade methanol (900 µL) and centrifuged at 13,000 x *g* for 10 min. Supernatants were dried and then suspended in 0.2 M TCA. TCA supernatants were then neutralized with 0.2 M NaOH and subjected to pre-column derivatization with o-phthaldialdehyde (OPA)/N-acetyl-L-cysteine (NAC) in 50% methanol. Diastereoisomer derivatives were resolved on a UHPLC Nexera X3 system (Shimadzu), using a Shim-pack GIST C18 reversed-phase column (3 μm, 4.0×160 mm, Shimadzu) under isocratic conditions (0.1 M sodium acetate buffer, pH 6.2, 1% tetrahydrofuran, and 1 mL/min flow rate). A washing step in 0.1 M sodium acetate buffer, 3% tetrahydrofuran, and 47% acetonitrile was performed after every run. Identification and quantification of d- and l-serine were based on retention times and peak areas, compared with external standards. CSF amino acid levels were expressed as µM.

For animal blood and tissue samples, serine enantiomers were quantified using a two-dimensional HPLC (2D-HPLC) system, as previously described(*41, 71*) with some modifications. Briefly, samples were homogenized in 19.5 volumes of distilled water (v/w). Homogenates were further mixed with 19 volumes methanol (v/v) (Fujifilm-Wako pure Chemical, Osaka, Japan) and centrifuged at 12,000 x *g* for 5 min. Supernatants were dried, re-suspended in 400 mM sodium borate, and derivatized with 4-fluoro-8-nitro-2,1,3-benzoxadiazole (NBD-F). NBD-conjugated amino acids were separated on an octadecylsilyl (ODS) column (Singularity RP18, 1.0 mm inner diameter (ID) x 250 mm) (designed by Kyushu University and KAGAI Inc., Japan) for separation in the first dimension, and enantiomers of amino acids were further separated using a Pirkle-type enantioselective column for separation in the second dimension (Singularity CSP-001S, 1.5-mm inner diameter (ID) x 250 mm) (designed by Kyushu University and KAGAMI Inc., Japan). Fluorescence intensity was detected at 530 nm with excitation at 470 nm.

### Analysis of pro- and anti-inflammatory cytokines in the CSF

Cytokines in CSF samples were analysed using a Bio-Plex multiplex cytokine assay (Bio-Rad Laboratories, Hercules, CA), according to the manufacturer’s instructions. Concentrations were calculated according to a standard curve generated for the specific target and expressed in pg/mL. All samples were analysed in triplicate. CSF molecules examined included: IL-1*β*, IL-2, IL-4, IL-5, IL-6, IL-7, IL-8, IL-9, IL-10, IL-12, IL-13, IL-15, IL17, Granulocyte-stem cell factor (G-SCF), Eotaxin, Fibroblast Growth Factor-Basic (FGF-Basic), monocyte chemoattractant protein 1 (MCP1), IP-10, Macrophage inflammatory protein-1*β* (MIP-1*β*), IL-1 receptor antagonist (ra), TNF, macrophage inflammatory protein 1-alpha (MIP-1*α*), MIP-1*β*, IFN-gamma, PDGF-BB, Rantes and VEGF.

### Animals

All animal experiments were performed in accordance with institutional guidelines of Keio University, and all protocols were approved by the Animal Experimental Committee. All animals had a C57BL6 background and were maintained in a specific pathogen-free environment. Dao-null mice (from homozygous G181R point mutations with C57BL6 background) were generated as previously described(*72*). Sr-KO mice were a gift from R. Konno(*47*). Dao-null/Sr-KO mice were produced by crossing Dao-null mice and Sr-KO mice. Vil1-Cre^+/-^/DAO^flox/flox^ mice were described previously(*42*). All animals were generated by *in vitro* fertilization and transplantation to surrogate mothers, and pups C-sectioned on E20 were fostered to CD-1 mothers to be colonized by microbiota of the same origin.

### EAE protocol and mouse tissue collection

Female mice with the C57BL6 background at eleven to thirteen weeks of age were used. MOG_35-55_ was emulsified in complete and incomplete Freund’s adjuvant (Chondrex, WA, USA) within 24 h before use and stored at 4°C until use. Mice were inoculated subcutaneously with the emulsion including 200 *μ*g MOG_35-55_, and injected intra-peritoneally with 300 ng of Pertussis toxin (PTX) in PBS, 2 and 24 h after the MOG inoculation. Animals were monitored daily from day 7 post-inoculation of MOG to assess development of paralysis. Motor functions were scored on scale 0 to 5.0 in 0.5 increments according to a protocol from Hooke Laboratories (MA, USA): 0 = normal; 1.0 = fully flaccid tail; 2.0 = weakness of hindlimbs; 3.0 = complete paralysis of hindlimbs; 4.0 = partial forelimb paralysis with hindlimb paralysis; 5.0 = moribund/death. When the clinical picture fell between two defined scores, intermediate scores were given. Mice with scores over 4.0 for two days were euthanized and scored as 5.0 for subsequent experiments. Control non-EAE mice were not inoculated with MOG or injected with PTX.

### Evaluation of BBB permeability *in vivo*

On day 11 (onset) after sensitization with MOG, mice were injected intravenously with 4 µL/g (BW) of 2% fluorescein-dextran (Merck, Darmstadt, Germany) in PBS into the tail vein. One hour after injection, under deep anesthesia, blood was drawn from inferior vena cava, and mice were perfused with ice-cold PBS. The cerebral cortex, cerebellum, and spinal cord were removed. Blood and tissues were further processed under protection from light. Plasma was isolated by centrifugation from blood to which 1 mg/mL 2Na-EDTA had been added. Tissues were homogenized in 5-fold v/w 1% Triton-X100 in PBS with metallic beads (ø = 3.5 mm) using a Micro Smash MS-100 (TOMY digital biology, Tokyo, Japan) at 3,500 rpm for 2 min, and centrifuged at 15,000 x *g* for 30 min at 4°C. Fluorescence of plasma and tissue supernatants was measured at 490/520 nm with a plate reader (Varioskan Lux, ThermoFisher, MA, USA) in a dark-wall, clear-bottom, 96-well plate. Fluorescence intensity in CNS tissue was standardized by that in plasma from the same mouse and reported as a percentage.

### Transmission electron microscopy

Deeply anaesthetized mice were perfused transcardially with ice-cold PBS and subsequently with 2% glutaraldehyde (EM grade, Electron Microscopy Science, PA, USA) in 0.1 M cacodylate buffer. Lumbar spinal cords were removed and post-fixed in 2% osmium tetra-oxide (Heraeus Chemicals South Africa, South Africa) for 2 h at 4°C. Then, tissues were dehydrated in a graded ethanol series (30 - 100%), embedded in epoxy resin, sliced at 80-90 nm using an ultramicrotome with diamond knives, and mounted on 200-mesh copper grids (EM fine grid, Nisshinn EM, Tokyo, Japan). Ultrathin sections were stained with uranyl acetate for 15 min and with lead staining solution for 2 min, and were visualized using an electron microscope (JEOL 1400Flash) at 100 kV.

### Statistical analysis

Statistical significance was determined with two-sided unpaired t-tests to compare two groups, or one-way analysis of variance (ANOVA) for multiple comparisons when data were normally distributed and had equal variance. If variances of the data were not equal, nonparametric tests were performed. Normality was tested using the Kolmogorov–Smirnov test. Data were shown as means with standard deviations or standard errors of the means, or as medians with interquartile ranges if not normally distributed. ROC analysis was performed using R studio and visualized using pROC and ggplot2 packages. The Youden index was used to determine cut-off values. Simple correlation was analyzed using Spearman’s correlation coefficient by R studio and visualized using corrplot packages. The Benjamini-Hockberg procedure was used to decrease the false discovery rate and to avoid false positives. To explore associations, after adjustment for possible confounding factors, i.e., age at lumbar puncture, disease duration, gender, BMI, OCB presence, or radiological disease activity, linear regression models were used. IBM SPSS Statistics for Windows (IBM Corp., Armonk, NY, USA) was used for statistical analyses of the clinical study, and Prism 9 (GraphPad Software) was for those of the animal study.

## List of Supplementary Materials

Materials and Methods

Figs. S1 to S12

Table S1

References 73-76

## Acknowledgments

We thank Alessia Casamassa, Mattia Miroballo, Giorgia Donati, Arianna De Rosa, Giulia Sansone, Giada Torresi, Takashi Sasaki, Maiko Nakane, Hiroshi Imoto, Tatsuhiko Ikeda, Eiichi Negishi, and Shoto Ishigo for their technical support; and Steven D Aird for editing the manuscript.

## Funding

NEXTGENERATIONEU (NGEU), the Ministry of University and Research (MUR), National Recovery and Resilience Plan (NRRP), Project MNESYS (PE0000006) (DN. 1553 11.10.2022) (AU, TN, RDV, DC)

PNRR-MCNT2-2023-12377850 (AU, DC)

Grant-in-Aid for Scientific Research (KAKENHI) 24K22061 (JS)

Keio Gijuku Fukuzawa Memorial Fund for Advancement of Education and Research (JS)

Keio Program for Promotion of Next Generation Research Projects Type A (JS)

## Author contributions

AU and JS conceived and coordinated the project. LG, MSB, RF, AF, and DC carried out the clinical evaluation of patients, collected the human CSF samples and analyzed the cytokine profile. KA, ST, XT, and JS performed animal experiments. RdV, TN, FE, KA, and MM carried out the HPLC experiments and data analysis. KA and RdV conducted molecular biological experiments and data analysis. KA and JS conducted transcriptomic analysis, and RR and AC performed the proteomic analysis. HM, JH, HO, and JN performed and analyzed animal MRI analysis. AH evaluated TEM images. AU, TK, MY, DC, and JS supervised the project. JS wrote the original manuscript, and AU and FE reviewed and edited it. All authors discussed the results and revised the manuscript.

## Data and materials availability

All data associated with this study are present in the paper or the Supplementary Materials.

