## Supplemental Methods and Figures for "Chiral shift toward D-serine reflects intrathecal inflammation in multiple sclerosis and counteracts motor impairment in a murine model"

### SUPPLEMENTARY MATERIALS AND METHODS

#### Western blotting

Spinal cord tissues were homogenized in lysis buffer (20 mM HEPES/NaOH pH 7.4, 150 mM NaCl, 1 mM EDTA, 0.5% Triton X-100, 1 mM DTT, and a protease inhibitor mixture (cOmplete; RochecO;Roche, Basel, Switzerland). Lysates (20 µg protein per lane) were subjected to SDS-PAGE, transferred to PVDF membranes, blocked with 10% skim-milk PBST, and blotted with a mouse monoclonal antibody to Sr (1:1000) (BD Biosciences, Franklin Lakes, NJ, USA), a rabbit polyclonal antibody to Dao (*72*), a mouse monoclonal antibody to β-actin (Sigma-Aldrich, Darmstadt, Germany), or rabbit monoclonal antibodies to Mbp, Gfap, Anx A2, or Gapdh (Cell Signaling Technology, Danvers, MA, USA), followed by appropriate HRP-conjugated antibodies. (ANXA2 info) Peroxidase activity was visualized with a western blot ECL substrate (Cytiva, Marlborough, MA, USA) and imaged with an ImageQuant LAS 4000mini (Cytiva).

#### RNA extraction and quantitative PCR

Total RNA was extracted from mouse brain regions using an RNeasy mini kit (QIAGEN) according to the manufacturer's instructions. RNA integrity was assessed by denaturing agarose gel electrophoresis (presence of sharp 28S, 18S, and 5S bands) and spectrophotometry (NanoDrop 2000, Thermo Scientific). Total RNA was purified to eliminate potential genomic DNA contamination using recombinant DNase (QIAGEN). 1 µg of total RNA of each sample was reverse-transcribed with QuantiTect Reverse Transcription (QIAGEN) using oligo-dT and a random primer mix according to the manufacturer's instructions. qPCR amplifications were performed using LightCycler 480 SYBR Green I Master (Roche Diagnostic) in a LightCycler480 Real Time thermocycler (Roche). The following protocol was used: 10 s for initial denaturation at 95°C followed by 40 cycles consisting of 10 s at 94°C for denaturation, 10 s at 60°C for annealing, and 6 s for elongation at 72°C. The following primers were used for mouse *Dao* forward, 5′-TTT TCT CCC GAC ACC TGG C-3′; and *Dao* reverse, 5′-TGA ACG GGG TGA ATC GAT CT-3′; and *Sr* forward, 5′-CCC TTG GTA GAT GCA CTG GT-3′ and *Sr* reverse, 5′-TCA GCA GCG TAT ACC TTC ACA C-3′. Relative transcription was evaluated using *Gapdh* as a housekeeping gene; *Gapdh* forward, 5′-CAT CAC TGC CAC CCA GAA GAC TG-3′, *Gapdh* reverse, 5′-ATG CCA GTG AGC TTC CCG TTC AG-3′.

#### RNA sequencing

Total RNA was extracted from lumber spinal cords as described above. *Only high-quality RNA preparations with RIN greater than 7.0 were used for RNA library* *construction.* A library was independently prepared from 1 μg of total RNA for each sample with an Illumina TruSeq Stranded mRNA Sample Prep Kit (Illumina, Inc., CA, USA). Poly-A-containing mRNA molecules, purified using poly-T-attached magnetic beads, were fragmented into small pieces with divalent cations at elevated temperature, and were copied into first strand cDNA using SuperScript II reverse transcriptase (Invitrogen) and random primers. This was followed by second strand cDNA synthesis using DNA Polymerase I, RNase H, and dUTP. These cDNA fragments went through an end-repair process, addition of a single 'A' base, and ligation of adapters. Products were then purified and enriched with PCR to create the final cDNA library. Libraries were quantified using KAPA Library Quantification kits for Illumina Sequencing platforms according to the qPCR Quantification Protocol Guide (KAPA Biosystems, Wilmington, MA, USA) and qualified using a TapeStation D1000 ScreenTape (Agilent Technologies, Santa Clara, CA, USA). Indexed libraries were then submitted to an Illumina NovaSeq (Illumina), and paired-end (2 x 100 bp) sequencing was performed by Macrogen Inc. (Seoul, South Korea).

Adapter sequences were trimmed from raw sequences with Trim Galore (https://www.bioinformatics.babraham.ac.uk/projects/trim_galore/) and trimmed sequences were mapped to the mouse genome (GRCm38/mm10) using HISAT2 (https://daehwankimlab.github.io/hisat2/). Aligned sequences were counted using FeatureCounts (https://subread.sourceforge.net/). Differentially expressed genes were identified with DESeq2 (https://bioconductor.org/packages/release/bioc/html/DESeq2.html). Gene Set Enrichment analysis was performed to find enriched biological pathways using MSigDB (https://www.gsea-msigdb.org/gsea/index.jsp) or ClusterProfiler (https://bioconductor.org/packages/release/bioc/html/clusterProfiler.html) with the following settings: pAdjustMethod = fd, pvalueCutoff = 0.05, minGSSize = 10, maxGSSize = 400 (*73, 74*).

**Proteomic analysis**

Spinal cord tissue samples were lysed in ice-cold lysis buffer (100 mM Tetraethylammonium bicarbonate TEAB, SDS 1%) and disrupted with two cycles of sonication at 20% amplitude for 30 sec on ice. Lysates were cleared by centrifugation at 16,000x g for 15 min at 4°C. Supernatants were transferred into new tubes and treated with 1 Unit of RQ1 DNase (Promega, Milan, Italy) for 1 h at room temperature. Protein concentration was determined using a Pierce BCA Protein assay kit (Thermo Scientific, Rodano MI, Italy). For each condition, equal amounts of proteins (100 μg in 100 μL of

100 mM TEAB) were reduced with 10 mM Tris-(2-carboxyethyl)-phosphine (TCEP) for 1 h at 55° C and alkylated with 18 mM iodoacetamide by incubating samples for 30 min at room temperature in the dark. Proteins were then precipitated overnight by adding six volumes of pre-chilled acetone. Following centrifugation at 8,000x g for 10 min at 4°C, protein pellets were resuspended in 100 µL of 100 mM TEAB and digested overnight with MS grade trypsin (Thermo Scientific, Rodano MI, Italy) at an enzyme/substrate ratio of 1:40 at 37° C. Resulting peptide mixtures were chemically labelled with TMT isobaric tags according to the manufacturer's instructions (Thermo Fisher Scientific). Peptides from biological triplicates of the 2 conditions were labeled using the following tags: 128N, 129N and 130N for EAE samples and 128C, 129C and 130C for EAE-D-serine samples. Briefly, 0.8 mg of TMT reagents in 41 µL of anhydrous acetonitrile were added to each sample. The reaction proceeded for 1 h and was then quenched for 15 min with hydroxylamine to a final concentration of 0.3%. The nine samples were then mixed in equal amounts and the resulting peptide mixture was fractionated using a Pierce™ High pH Reversed-PhasePeptide Fractionation Kit, following the manufacturer's instructions (Thermo Fisher Scientific). Briefly, labelled peptides were loaded onto an equilibrated, high-pH, reversed-phase fractionation spin column. Peptides bound to the hydrophobic resin under aqueous conditions and were desalted by washing the column with water during low-speed centrifugation. A step gradient of increasing acetonitrile concentrations in a volatile high pH elution solution was then applied to the columns to elute bound peptides into eight fractions collected by centrifugation. Each fraction was dried in a vacuum centrifuge and re-suspended in 20 µL of 0.1 % formic acid.

The eight fractions were analyzed with high-resolution nanoLC-Tandem Mass Spectrometry using a Q-Exactive Orbitrap mass spectrometer equipped with an EASY-Spray nano-electrospray ion source (Thermo Fisher Scientific, Rodano MI, Italy) and coupled to a Thermo Scientific Dionex UltiMate 3000RSLC nano system (Thermo Fisher Scientific) (Di Matteo et al. 2020). Solvent composition was 0.1% formic acid in water (solvent A) and 0.1% formic acid in acetonitrile (solvent B). Peptides were loaded on a trapping PepMap™100 Cartridge Column $C_{18}$ (300 µm x 0.5 cm, 5 µm,100 Å) and desalted with solvent A for 3 min at a flow rate of 10 µL/min. After trapping, eluted peptides were separated on an EASY-Spray analytical column (50 cm x 75 µm ID PepMap RSLC C18,3 µm, 100 Å), heated at 35°C at 300 nL/min, applying the following gradient: 5% B for 3 min, from 5% to 27.5% B in 222 min, from 27.5% to 40% B in 10 min, from 40% to 95% B in 1 min. Washing (95% B for 4 min) and re-equilibration (5% B for 24 min) steps were always included at the end of the gradient. Eluting peptides were analyzed on the Q-Exactive mass spectrometer operating in positive polarity mode with a capillary temperature of 280°C and a potential of 1.9 kV applied to the capillary probe. Full MS survey scan resolution was set to 70,000 with an automatic gain control (AGC) target value of 3 x $10^6$ for a scan range of 375-1500 m/z and maximum ion injection time (IT) of 60 ms. The mass (m/z) 445.12003 was used as a lock mass. A data-dependent top 12 method was operated, during which high-energy collisional dissociation (HCD) spectra were obtained at 35,000 MS2 resolution with an AGC target of 1 x $10^5$ for a scan range of 200-2000 m/z, maximum IT of 120 ms, 1.6 m/z isolation width and normalized collisional energy (NCE) of 32. Precursor ions targeted for HCD were dynamically excluded for 30s. Full scans and Orbitrap MS/MS scans were acquired in profile mode, whereas ion trap mass spectra were acquired in centroid mode. Charge state recognition was enabled by excluding unassigned and 1, 7, 8, >8 charged states. All data were acquired with the Xcalibur 3.1 software (Thermo-Fisher Scientific).

For data processing, acquired raw files were analyzed with Thermo Scientific Proteome Discoverer 2.4 software (Thermo Fisher Scientific) using the SEQUEST HT search engine. HCD MS/MS spectra were searched against the *Mus musculus* database (version 03-2023, number of entries 37,654 sequences) assuming trypsin (Full) as digestion enzyme and allowing two missed cleavages. Mass tolerances were set to 10 ppm and 0.02 Da for precursor and fragment ions, respectively. Oxidation of methionine (+15.995 Da) was set as a dynamic modification. Carbamidomethylation of cysteine (+57.021 Da) and the TMT label on lysines and the N-terminus (229.1629) were set as static modifications. False discovery rates (FDRs) for peptide spectral matches (PSMs) were calculated and filtered using the Percolator node in Proteome Discoverer, with the following settings: Maximum Delta Cn 0.05, a strict target FDR of 0.01, a relaxed target FDR of 0.05 and validation based on q-value. Protein identifications were accepted when the protein FDR was below 1% and when proteins were identified with at least two peptides.

For bioinformatic analyses, proteins with log2 fold change values (log2FC) ≥ 0.3 and ≤ 0.3 were considered as differentially expressed (DE). Enriched Gene Ontology (GO) Molecular function terms for DE proteins were extracted using a module integrating the ClusterProfiler R package (*75*). Plots were visualized using the online platform for data analysis and visualization available at https://www.bioinformatics.com.cn.

#### *q*-Space diffusion MRI

An MR scanner equipped with a 9.4 T magnet with high gradient strength (660 mT/m) (Biospec 94/30; Bruker, Billerica, MA, USA) was used. Solenoid coils with inner diameters (ID) of 28 mm tuned to 400 MHz for proton resonance were used for measurements. Spinal cords extracted from WT, Sr-KO, and Dao-null mice were inserted into a 20-mm ID acrylic tube containing electronic liquid (Fluorinert FC-72; 3M, Maplewood, MA, USA) and were immediately subjected to imaging. For the pulse-field gradient spin-echo (PGSE) sequence, the following parameters were used: repetition time (TR) = 2000 ms, echo time (TE) = 32 ms, matrix = 225 x 300, field of view (FOV) = 18 x 24 mm2, and slice thickness 1 mm and 12 motion-probing axes with nine b-value steps (0-10,000 s/mm$^2$) for measurements. A *q*-space myelin map was obtained, as previously reported (*76*).

### SUPPLEMENTARY FIGURES

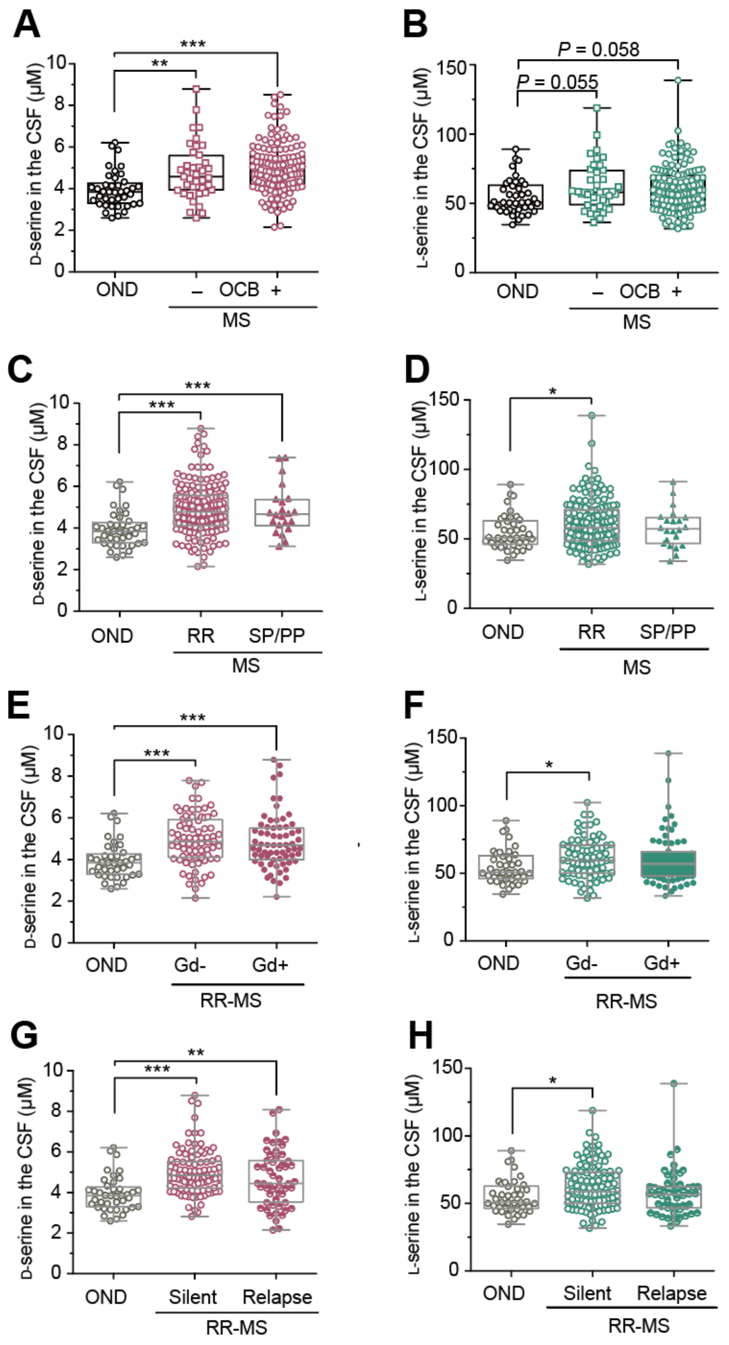


**Fig. S1. Serine enantiomers in the CSF of patients with MS.** (**A-H**) Levels of D-(A, C, E, and G) or L-serine (B, D, F, and H) in the CSF drawn from patients with OND (n = 40) and MS (n = 179) at diagnosis were quantified with HPLC. Samples from patients with MS were categorized with detectability of OCB (A, B) or disease types (C, D). Those with RR-MS were categorized with Gd extravasation in MRI (E, F) or disease activity (G, H). Data are shown in box plots. ****P* < 0.001, ***P* < 0.01, **P* < 0.05; Mann-Whitney test.

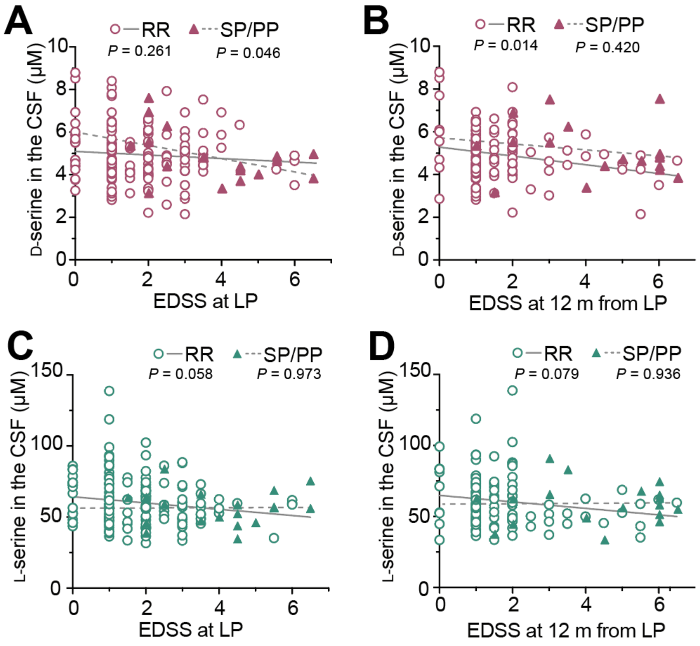


**Fig. S2. Serine enantiomers in the CSF and EDSS scores of patients with MS.** (**A-D**) Correlations between CSF levels of D-(A and B) or L-serine (C and D) drawn from patients with MS (n = 179) at diagnosis and EDSS scores of patients with RR- (n = 157) or SP/PP-MS (n = 22) at lumber puncture (LP) (A and C) or at 12 months after LP (C and D) were analyzed. *P*-values, linear regression analysis.

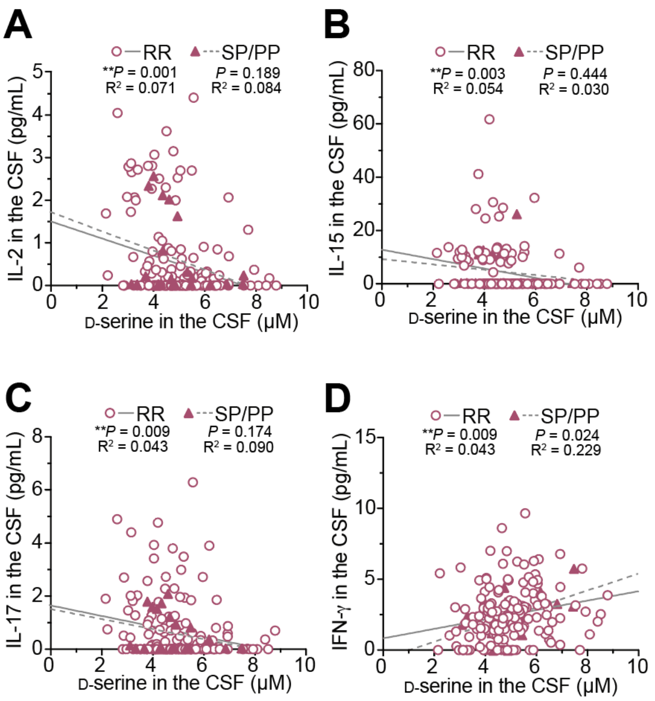


**Fig. S3. Associations between D-serine and cytokines in the CSF of patients with MS.** (**A-D**) Correlations between CSF levels of D-serine and IL-2, IL-15, IL-17, and IFN-g in patients with RR- (n = 157) or SP/PP-MS (n = 22) were analyzed. *P*-values, linear regression analysis.

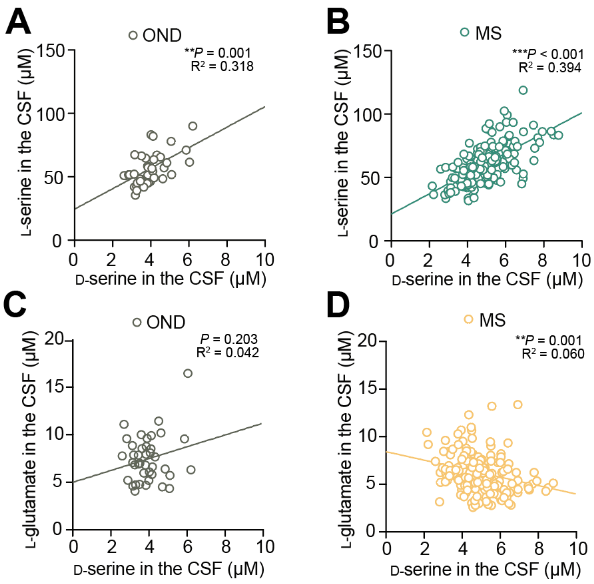


**Fig. S4. Associations of D-serine and L-serine/L-glutamate in CSF with MS. (A-D)** Correlations between CSF D-serine and L-serine in patients with OND (n = 40, A) or MS (n = 179, B) or between CSF D-serine and L-glutamate in patients with OND (C) or MS (D) are shown. *P*-values, linear regression analysis.

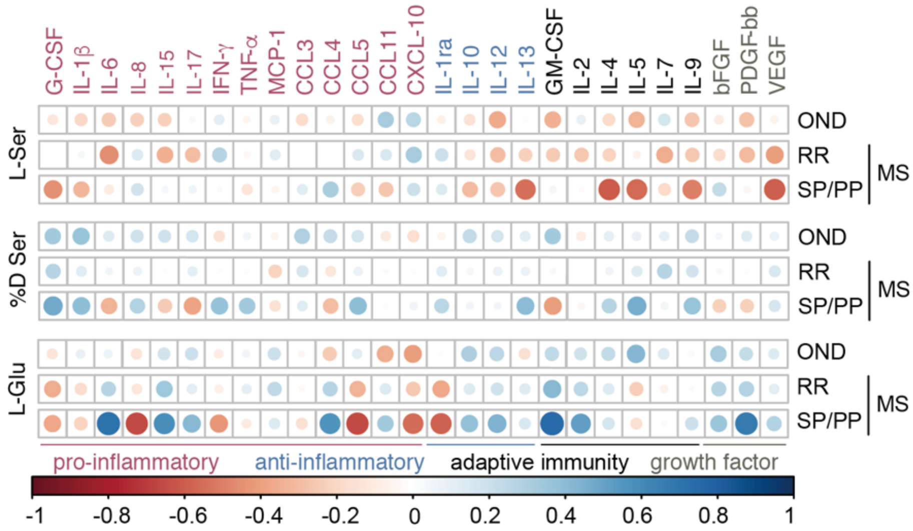


**Fig. S5. Correlations between CSF amino acids and cytokines in CSF with MS.** Heatmaps show Spearman's correlation coefficients between CSF L-serine, %D serine, and L-glutamate and cytokines in CSF of patients with OND (n = 40) and RR- (n = 157) or SP/PP-MS (n = 22).

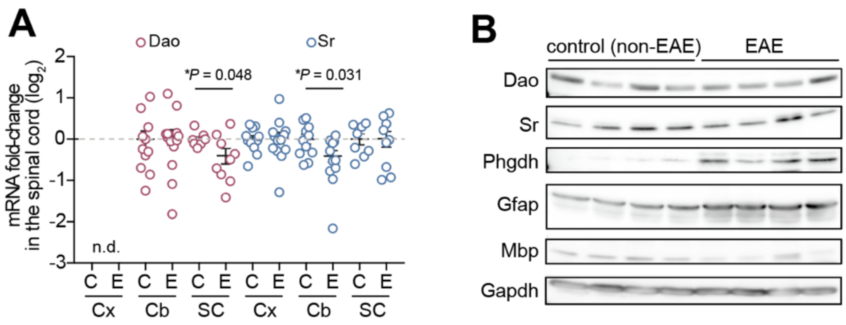


**Fig. S6. Transcription and protein expression of metabolic enzymes for serine enantiomers in mice with EAE.** (**A**) Transcription of Dao and Sr genes in the cerebral cortex (Cx), cerebellum (Cb), and spinal cord (SC) in EAE mice (E) and controls (C) were evaluated with qPCR. Data are shown as mean ± s.e.m. *$P < 0.05$; unpaired two-tailed t-test. (**B**) Protein expression of Dao, Sr, Phgdh, Gfap, Mbp, and Gapdh in spinal cords from mice with EAE or controls were tested with western blotting.

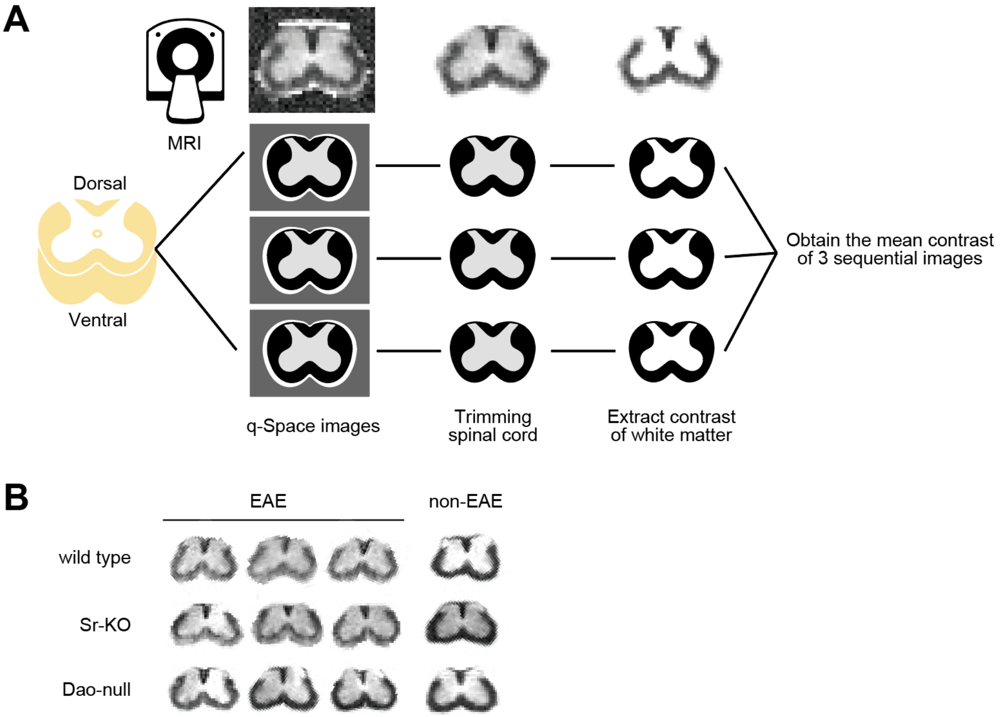


**Fig. S7. Impact of D-serine metabolism on myelin imaging in spinal cords of mice with EAE.** (**A**) Lumbar spinal cords were isolated from mice, fixed, and processed for MRI. Image contrast was obtained from *q*-space diffusion images of MRI after trimming images of spinal cords and extracting their white matter. (**B**) Representative trimmed images of spinal cords from wild-type, Sr-KO, and Dao-null mice with or without EAE were obtained using *q*-space diffusion MRI.

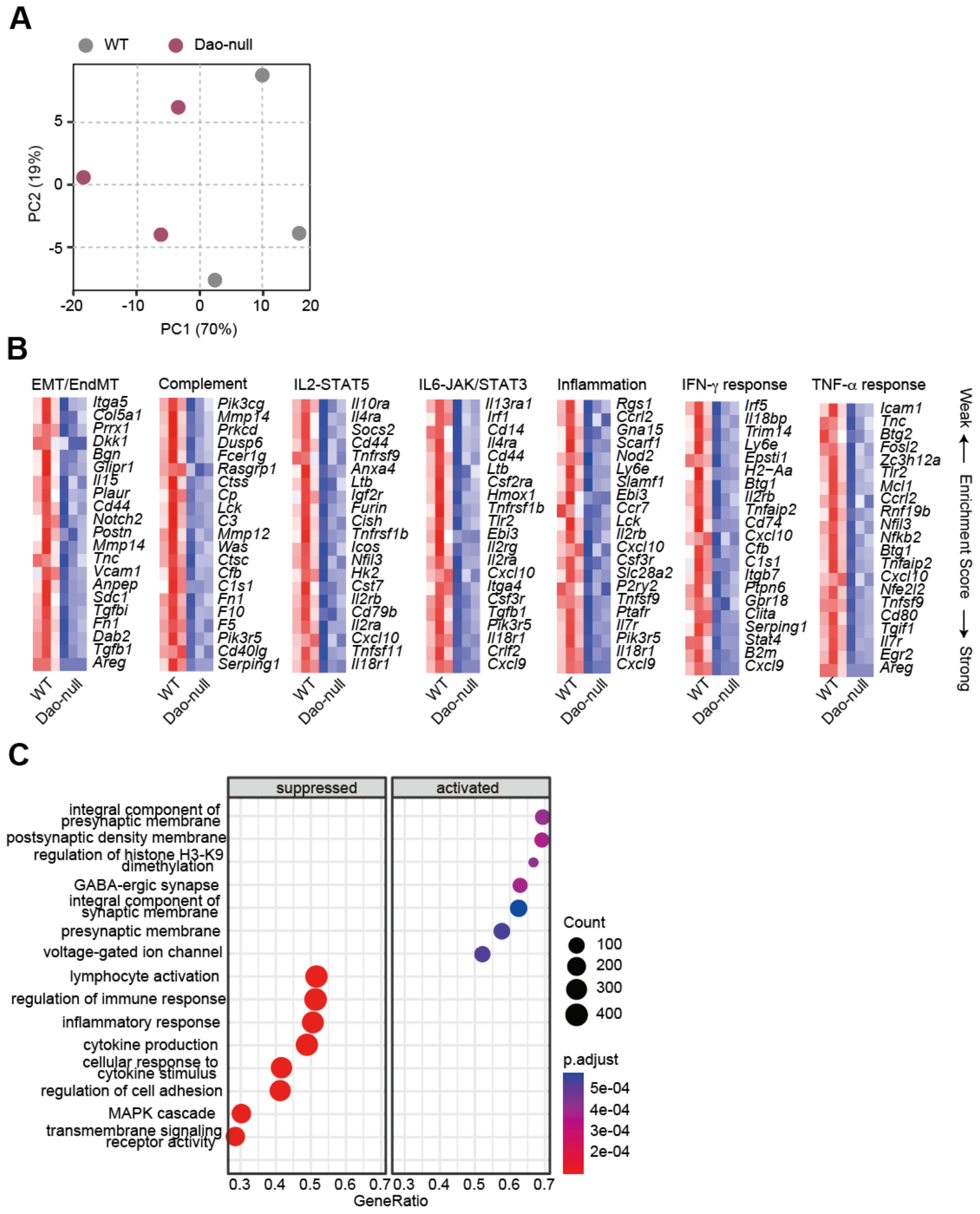


**Fig. S8. Transcriptomic changes by loss of Dao in spinal cords of mice with EAE.** (A-C) RNA-seq was performed using spinal cords from Dao-null and WT mice with EAE at an acute phase (18 days after MOG injection). A PCA plot of RNA-seq results (A). Heat maps showing z-scores of gene transcription categorized by the presented gene ontology significantly altered by loss of Dao analyzed by GSEA using MSigDB (B). A dotplot indicating activated/suppressed gene ontology analyzed by GSEA with ClusterProfiler (C).

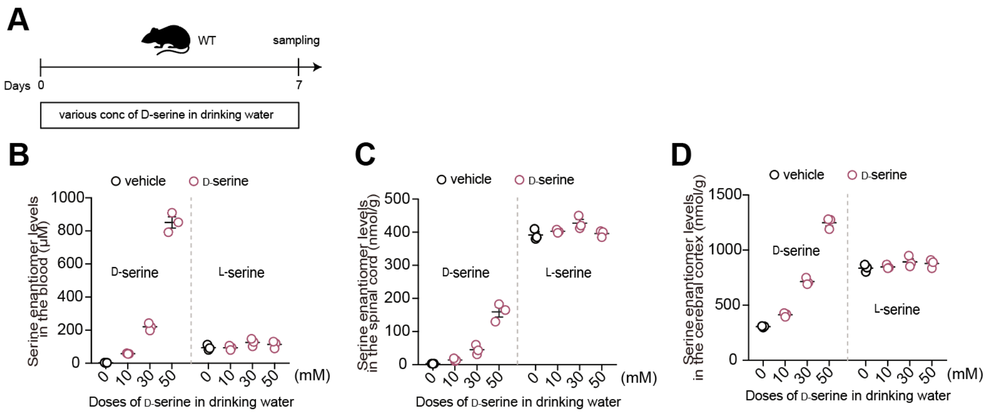


**Fig. S9. Concentrations of serine enantiomers in mice after loading D-serine in the drinking water.** (**A**) Mice ingested water containing D-serine at 0, 10, 30, or 50 mM ad libitum for 7 days, and were sacrificed for sampling (n = 3, each group). (**B**-**D**) Levels of serine enantiomers in plasma (B), spinal cords (C), and cerebral cortex (D) of these mice were quantified using the 2D-HPLC. Data are shown as means ± s.e.m.

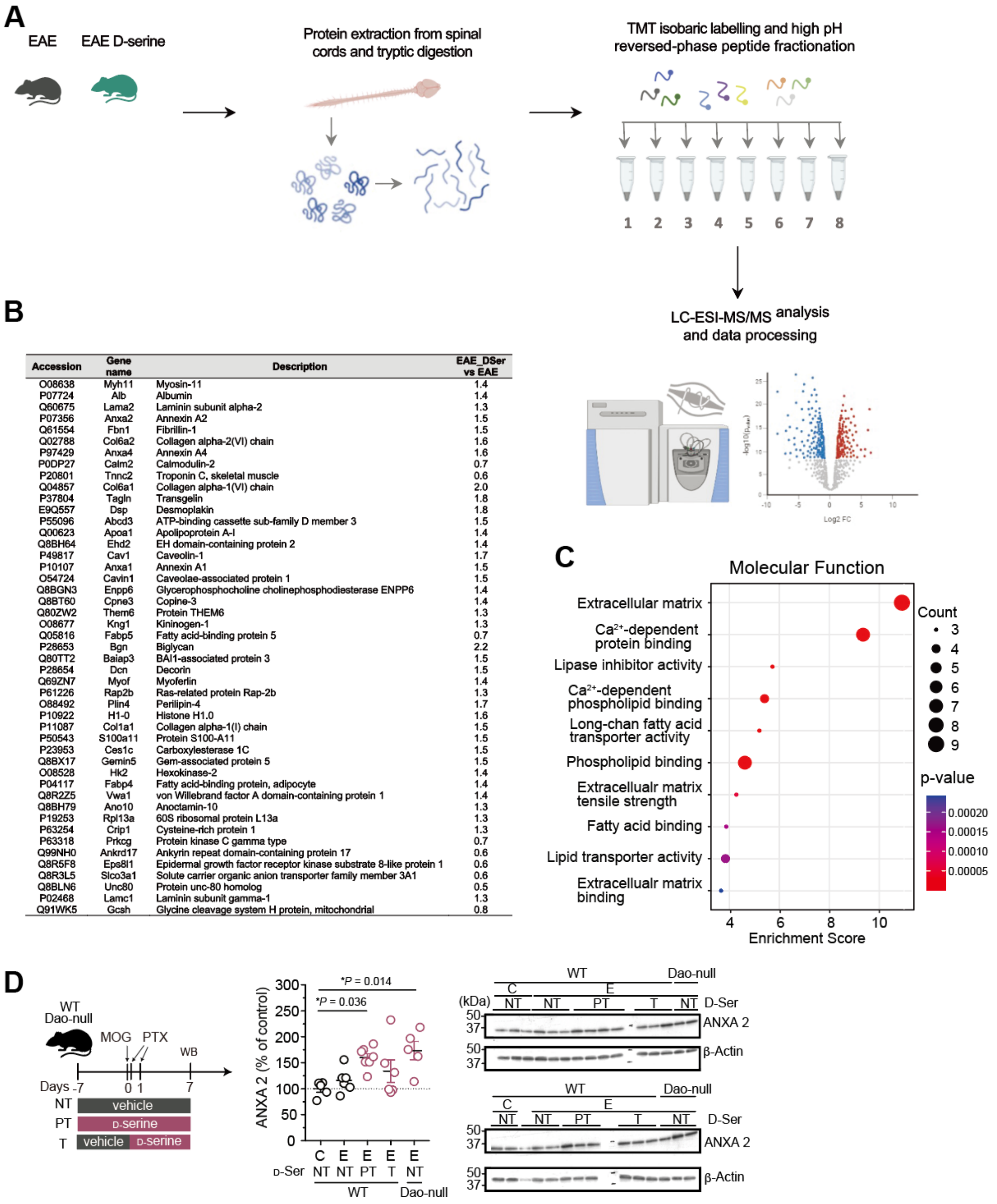

| Accession | Gene name | Description | EAE_DSer vs EAE |
|---|---|---|---|
| O08638 | Myh11 | Myosin-11 | 1.4 |
| P07724 | Alb | Albumin | 1.4 |
| Q60675 | Lama2 | Laminin subunit alpha-2 | 1.3 |
| P07356 | Anxa2 | Annexin A2 | 1.5 |
| Q61554 | Fbn1 | Fibrillin-1 | 1.5 |
| Q02788 | Col6a2 | Collagen alpha-2(VI) chain | 1.6 |
| P97429 | Anxa4 | Annexin A4 | 1.6 |
| P0DP27 | Calm2 | Calmodulin-2 | 0.7 |
| P20801 | Tnnc2 | Troponin C, skeletal muscle | 0.6 |
| Q04857 | Col6a1 | Collagen alpha-1(VI) chain | 2.0 |
| P37804 | Tagln | Transgelin | 1.8 |
| E9Q557 | Dsp | Desmoplakin | 1.8 |
| P55096 | Abcd3 | ATP-binding cassette sub-family D member 3 | 1.5 |
| Q00623 | Apoa1 | Apolipoprotein A-I | 1.4 |
| Q8BH64 | Ehd2 | EH domain-containing protein 2 | 1.4 |
| P49817 | Cav1 | Caveolin-1 | 1.7 |
| P10107 | Anxa1 | Annexin A1 | 1.5 |
| O54724 | Cavin1 | Caveolae-associated protein 1 | 1.5 |
| Q8BGN3 | Enpp6 | Glycerophosphocholine cholinephosphodiesterase ENPP6 | 1.4 |
| Q8BT60 | Cpne3 | Copine-3 | 1.4 |
| Q80ZW2 | Them6 | Protein THEM6 | 1.3 |
| O08677 | Kng1 | Kininogen-1 | 1.3 |
| Q05816 | Fabp5 | Fatty acid-binding protein 5 | 0.7 |
| P28653 | Bgn | Biglycan | 2.2 |
| Q80TT2 | Baiap3 | BAI1-associated protein 3 | 1.5 |
| P28654 | Dcn | Decorin | 1.5 |
| Q69ZN7 | Myof | Myoferlin | 1.4 |
| P61226 | Rap2b | Ras-related protein Rap-2b | 1.3 |
| O88492 | Plin4 | Perilipin-4 | 1.7 |
| P10922 | H1-0 | Histone H1.0 | 1.6 |
| P11087 | Col1a1 | Collagen alpha-1(I) chain | 1.5 |
| P50543 | S100a11 | Protein S100-A11 | 1.5 |
| P23953 | Ces1c | Carboxylesterase 1C | 1.5 |
| Q8BX17 | Gemin5 | Gem-associated protein 5 | 1.5 |
| O08528 | Hk2 | Hexokinase-2 | 1.4 |
| P04117 | Fabp4 | Fatty acid-binding protein, adipocyte | 1.4 |
| Q8R2Z5 | Vwa1 | von Willebrand factor A domain-containing protein 1 | 1.4 |
| Q8BH79 | Ano10 | Anoctamin-10 | 1.3 |
| P19253 | Rpl13a | 60S ribosomal protein L13a | 1.3 |
| P63254 | Crip1 | Cysteine-rich protein 1 | 1.3 |
| P63318 | Prkcg | Protein kinase C gamma type | 0.7 |
| Q99NH0 | Ankrd17 | Ankyrin repeat domain-containing protein 17 | 0.6 |
| Q8R5F8 | Eps8l1 | Epidermal growth factor receptor kinase substrate 8-like protein 1 | 0.6 |
| Q8R3L5 | Slco3a1 | Solute carrier organic anion transporter family member 3A1 | 0.6 |
| Q8BLN6 | Unc80 | Protein unc-80 homolog | 0.5 |
| P02468 | Lamc1 | Laminin subunit gamma-1 | 1.3 |
| Q91WK5 | Gcsh | Glycine cleavage system H protein, mitochondrial | 0.8 |



**Fig. S10. Impact of D-serine on proteomes of spinal cords in EAE mice.** (**A**) Overview of a strategy based on tandem mass tags (TMT) profiling for in-depth quantification of spinal cord proteome by high-resolution mass spectrometry. Mice with/without D-serine in the drinking water before and after MOG injection were used. At a pre-symptomatic stage (7 days after MOG injection), spinal cords were dissected from the mice and processed for proteome analysis. (**B**) A list of significantly up- or down-regulated proteins in spinal cords of EAE mice treated with D-serine compared with those treated with vehicle (ratios of protein expression shown as D-serine/vehicle). (**C**) Enriched gene ontology (GO) terms of differentially expressed proteins in spinal cords from EAE mice treated with D-serine vs vehicle. Color of the plots shows *P*-values and their sizes represent protein counts. (**D**) Western blotting of ANXA2 and b-Actin was performed using spinal cords from WT or Dao-null mice with EAE (E) or non-EAE (C) treated with vehicle and/or D-serine for the indicated period at day 7 after MOG induction. Vehicle only (NT), D-serine before and after MOG (PT), and D-serine after MOG only (T). ANXA2 expression was standardized against that of b-Actin and shown as relative values compared to non-EAE mice treated with vehicle (WT/C/NT, n = 5; WT/E/NT, n = 6; WT/E/PT, n = 7; WT/E/T, n = 6; and Dao-null/E/NT, n = 5)(middle panel). Data are shown as means ± s.e.m. *$P < 0.05$; one-way ANOVA with Tukey's test.

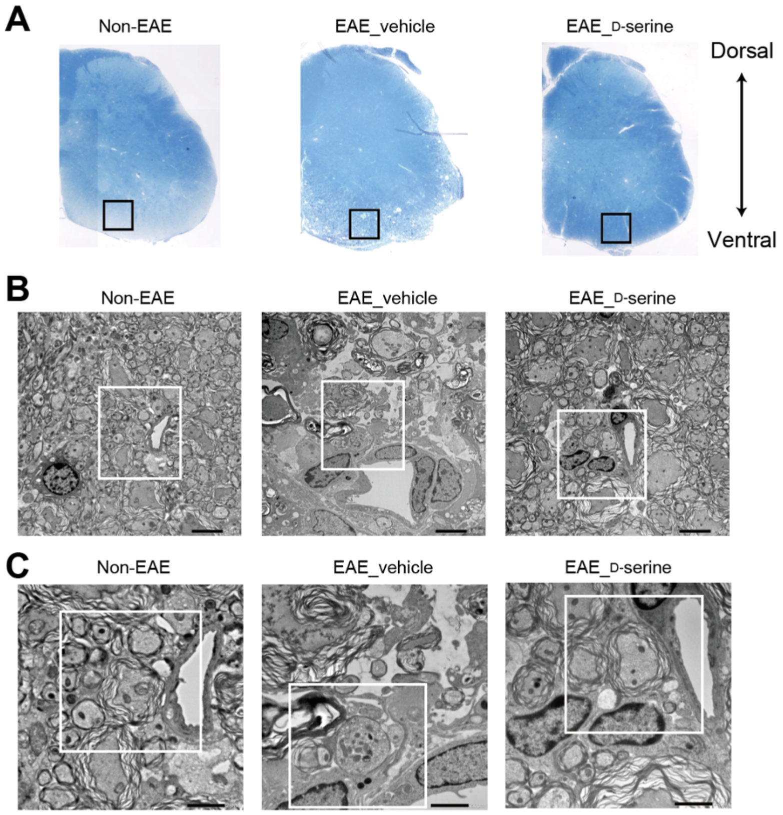


**Fig. S11. Effect of D-serine on myelin and vascular pathology of EAE.** Lumber spinal cords from non-EAE or EAE mice treated with D-serine or vehicle at an acute phase (15 days after MOG injection) were processed for transmission electron microscopy (TEM). (**A**) Light microscopic images show horizontal sections of spinal cords stained with toluidine blue. Squared regions were trimmed for electron microscopy. (**B** and **C**) Representative TEM images indicate enlarged anterior funiculus of the white matter in spinal cords (x 1000, B; x 3000, C). Whereas myelin sheath structure was extensively destroyed (demyelination) and bare axons were dispersed in the EAE spinal cords treated with vehicle (B and C, middle), axons enwrapped with myelin remained intact in EAE spinal cords treated with D-serine (B and C, right). In EAE spinal cord treated with vehicle, vascular endothelial cells were swollen and perivascular structures were damaged (B and C, middle). In contrast, basement membrane structure was preserved in EAE spinal cord treated with D-serine (B and C, right). Error bars, 5 µm (B) and 2 µm (C). Squared regions in (B) were enlarged in (C) and those in (C) were magnified in Fig. 5E.

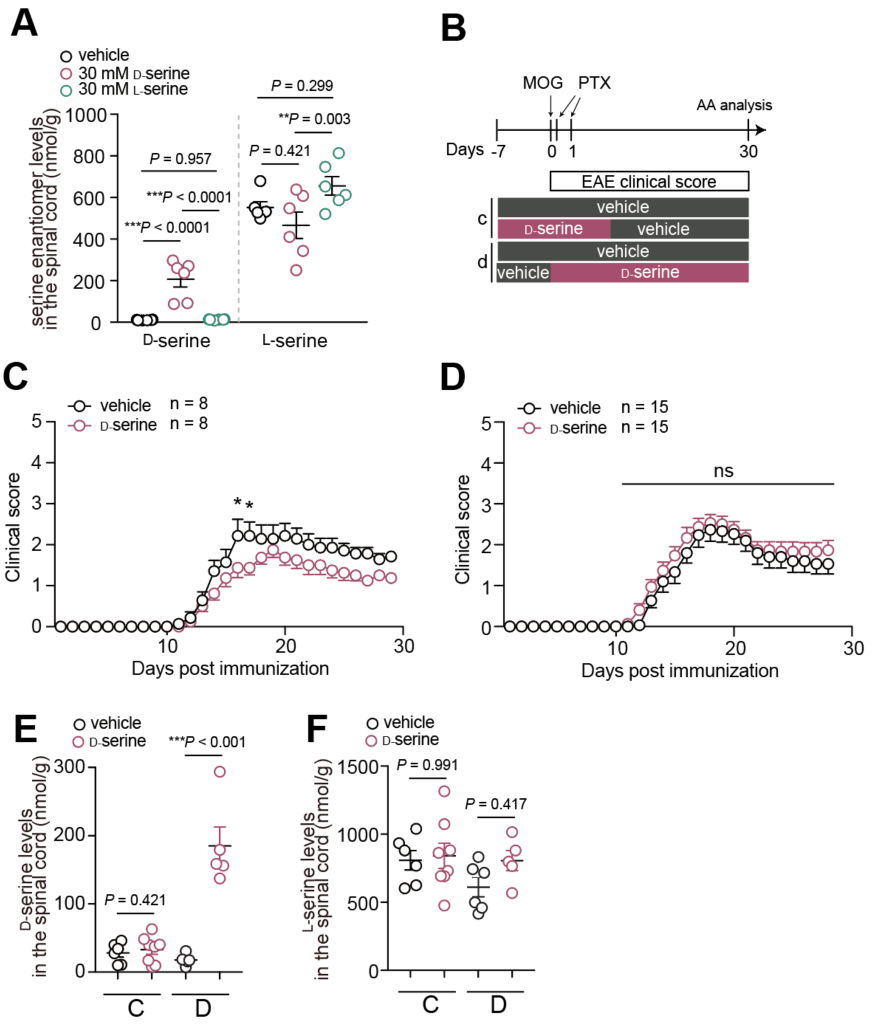


**Fig. S12. Difference in EAE improvement by D-serine depending on timing of its administration.** (**A**) Serine enantiomers in spinal cords from EAE mice treated with vehicle, D-serine, or L-serine were quantified at a chronic phase (30 days after MOG) using the 2D-HPLC (n = 6, each). (**B-F**) A scheme indicates the timing of D-serine treatment in EAE mice (B). C57BL6 mice were treated with 30 mM D-serine in the drinking water for 14 days before and after MOG injection (C, E, F) or for 30 days after MOG injection (D, E, F). Clinical scores of EAE mice treated with D-serine or vehicle at different timing were evaluated after MOG injection (n = 8, each, C; and n = 15, each, D). D- (E) or L-serine (F) in the spinal cords of EAE mice treated with D-serine or vehicle at different timing was quantified at 30 days after MOG injection using the 2D-HPLC (n = 6, each). Data are shown as means ± s.e.m. ***$P < 0.001$, **$P < 0.01$, *$P < 0.05$; one-way ANOVA with Tukey's test (A), Mann-Whitney test (E and F), and two-way ANOVA with Sidak's test (C and D).
