## Supplemental Table for "Chiral shift toward D-serine reflects intrathecal inflammation in multiple sclerosis and counteracts motor impairment in a murine model"

**Table S1. Demographic and clinical characteristics of multiple sclerosis and other neurological diseases subjects.**

| | | ***Multiple Sclerosis*** | | |
|---|---|---|---|---|
| *Demographic and clinical characteristics* | ***OND** (n=40)* *median (min; max)* | ***RR-MS** (n=157)* *median (min; max)* | ***SP/PP-MS** (n=22)* *median (min; max)* | ***Total MS** (n=179)* *median (min; max)* |
| Sex (Female / Male) | 24 / 16 | 109 / 48 | 9 / 13 | 118 / 61 |
| Age at LP (years) | 41.76 (16.91; 64.86) | 35.61 (16.30; 75.55) | 48.62 (32.73; 72.95) | 37.99 (16.3;75.55) |
| BMI | 23.89 (18.49; 39.1) *n=37* | 24.44 (17.51; 39.58) *n=136* | 26.21 (16.98; 42.28) *n=20* | 24.67 (16.98; 42.28) *n=156* |
| Radiological activity ($Gd^-$ / $Gd^+$) | 9 / 1 | 75 / 67 | 16 / 5 | 91 / 72 |
| OCB presence (No / Yes) | 22 / 2 | 37 /114 | 2 / 20 | 39 / 134 |
| MS disease duration (months) | - | 6.98 (0; 371.53) *n=150* | 20.57 (6.33; 175.40) | 9.83 (0.06;371.53) *n=172* |
| EDSS score at LP | - | 1.50 (0; 6.00) | 4.25 (1.50; 6.50) *n=20* | 2 (0; 6.50) *n=177* |
| EDSS score at 1 year from LP | - | 1 (0; 6.50) *n=118* | 4.75 (1; 6.50) *n=18* | 1.50 (0; 6.50) *n=136* |
| CSF lactate (mmol/l) | 1.40 (1.00; 3.00) *n=39* | 1.50 (0.90; 2.20) *n=153* | 1.55 (1.20; 2.10) | 1.50 (0.90; 2.20) *n=175* |

Abbreviations: Lumbar puncture (LP), gadolinium (Gd), Oligoclonal bands (OCB), body max index (BMI), Expanded Disability Status Scale (EDSS), cerebrospinal fluid (CSF), other neurological diseases (OND), multiple sclerosis (MS), primary progressive (PP), relapsing-remitting (RR), secondary progressive (SP).
